# *In vivo* anti-tumor activity of high-dose parenteral ascorbic acid is mediated primarily via cofactor activity, not via oxidative stress

**DOI:** 10.1101/2025.05.21.655101

**Authors:** Talia Akram, Rebecca A Luchtel, Vinay Dubey, Soma Seal, Ritesh Aggarwal, Hiroaki Sai, Niraj K Shenoy

**Affiliations:** Department of Medicine, Feinberg School of Medicine, Northwestern University, Chicago, IL 60611; Department of Medicine, Albert Einstein College of Medicine, Bronx, NY 10461; Analytical BioNanoTechnology Equipment Core (ANTEC), Simpson Querrey Institute, Northwestern University, Chicago, IL 60611; Department of Pathology, Feinberg School of Medicine, Northwestern University, Chicago, IL 60611; Robert H. Lurie Comprehensive Cancer Center, Northwestern University, Chicago, IL 60611

## Abstract

The anti-tumor effect of high-dose ascorbic acid (AA) has been demonstrated in multiple *in vitro* and *in vivo* cancer models with the postulation of two primary categories of mechanisms: antioxidant/cofactor activity and H_2_O_2_-mediated oxidative damage. Both mechanisms have been conclusively demonstrated *in vitro*. However, while parenteral high-dose AA-induced cofactor activity (TET-mediated DNA demethylation and prolyl/asparaginyl hydroxylase-mediated HIF activity inhibition via reduction of enzymatic Fe^3+^ to Fe^2+^) has been demonstrated intratumorally *in vivo* in multiple models, the cumulative data on parenteral high-dose AA-induced intratumoral oxidative damage *in vivo* has been inconclusive. Furthermore, the relative contribution of the seemingly opposing mechanisms towards *in vivo* anti-cancer activity has not been studied concurrently. We therefore sought to definitively delineate the roles of both antioxidant/cofactor activity and prooxidant functions of high-dose AA in the *in vivo* anti-tumor response. Using two syngeneic mouse tumor models, the AA-sensitive A20 model and the AA-resistant Renca model, we assessed markers of DNA and lipid oxidative damage as well as the specific roles of TET2 and AA transporter SLC23A2 in the anti-tumor response to parenteral high-dose AA. In the sensitive A20 model, loss of either *Tet2* or *Slc23a2* fully reversed anti-tumor activity. Similarly, overexpression of *Tet2* in the resistant Renca model (which expresses high baseline levels of AA transporters SLC23A1 and SLC23A2, but does not express TET2), resulted in increased CD8^+^T cell infiltration and dramatic reduction in tumor growth overall. In both A20 and Renca models, high-dose parenteral AA *increased* total intratumoral antioxidant capacity, and this was attenuated by *Slc23a2* knockdown in A20. High-dose AA treatment also resulted in a *Tet2*- and *Slc23a2*-dependent increase in intratumoral 5-hydroxymethylcytosine. Intracellular oxidative damage markers, 8-OHdG and 4-HNE, were not induced in tumors by high-dose AA in either model. In contrast, these markers were robustly induced *in vitro* by high-dose AA in A20 and Renca cells. Using dynamic real-time extracellular H_2_O_2_ measurements with high-dose AA, difference in molecular oxygen concentration between standard *in vitro* and hypoxic *in vivo* conditions was identified as an important factor underlying the marked discrepancy between the abundant *in vitro* and absent *in vivo* intratumoral oxidative stress with high-dose AA. Furthermore, using additional syngeneic models resistant (MB49) and sensitive (MC38) to AA-induced potentiation of anti-PD1 checkpoint inhibition, we demonstrate that very low catalase expression does not confer sensitivity to high-dose AA *in vivo* (further arguing against the H_2_O_2_ mechanism *in vivo*), that TET2 expression alone is not sufficient to drive an AA-induced anti-tumor response (either as a single agent or in combination with immunotherapy), and that high-dose AA can significantly enhance the efficacy of anti-PD1 immunotherapy even in the absence of single-agent activity. Our data strongly indicate that the *in vivo* anti-tumor effect of high-dose parenteral AA-including potentiation of immunotherapy-is mediated primarily by its specific antioxidant/cofactor activity (with TET2 expression likely being necessary but certainly not sufficient), and not via oxidative stress. Collectively, the study represents a paradigm shift in our understanding of the cumulative mechanisms of *in vivo* anti-cancer activity of high-dose AA, with critical implications not just for the clinical translation of AA as an anti-cancer agent (including in enhancing immunotherapy efficacy) but also the field of free radical biology.

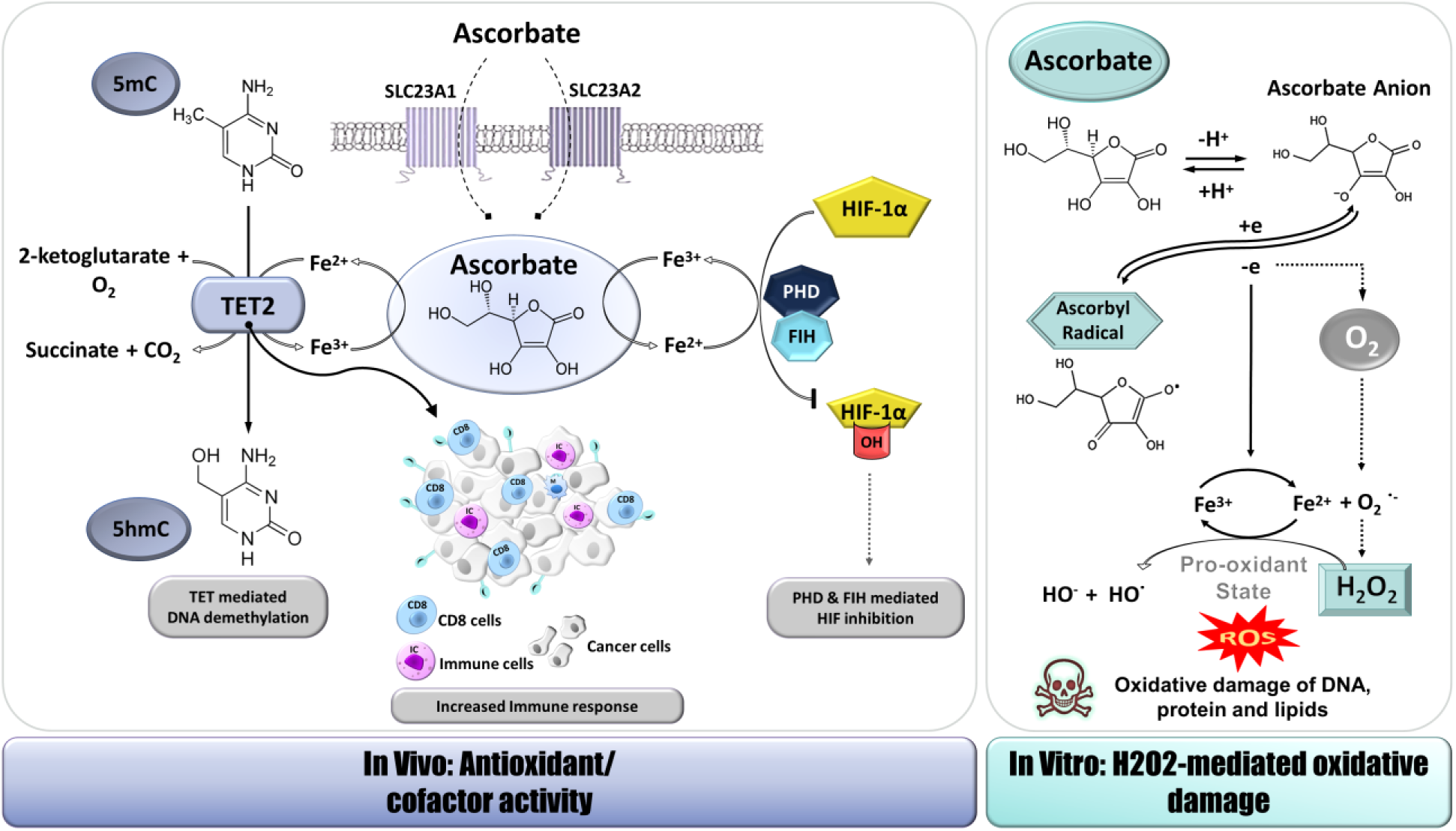

## Background

There has been a renewed interest in high-dose parenteral ascorbic acid as an anti-cancer agent in recent years owing to increased understanding of biologic mechanisms, pharmacokinetics, and demonstration of anti-tumor activity in various preclinical models, including enhancement of immunotherapy efficacy ^1–8^. Two categories of mechanisms have gained prominence based on preclinical studies: co-factor activity (TET-mediated demethylation and HIF downregulation) and H_2_O_2_-mediated oxidative stress. Although these two categories of mechanisms have been studied separately in various models, they are seemingly opposing (antioxidant/cofactor vs prooxidant effects) and have not been *concurrently and definitively* assessed in animal models to determine their relative contribution to parenteral high-dose ascorbate’s *in vivo* anti-cancer effects.

It has been proposed that high-dose parenteral ascorbate acts as a prooxidant *in vivo*, resulting in ascorbate radical and H_2_O_2_ generation, similar to the indisputable *in vitro* oxidative effects with high-dose AA ^9,10^. In support of this, high-dose ascorbic acid (4g/kg) administered intraperitoneally resulted in ⁓150uM H_2_O_2_ in the probe eluate from the tumor interstitial space of a xenograft mouse model, for 3 hours after administration ^11^. However, data on the intratumoral intracellular effects of H_2_O_2_ production, i.e. oxidative damage of DNA and/or lipid/protein oxidation leading to growth inhibition, following administration of high-dose AA intraperitoneally or intravenously, has been inconclusive ^7,12^.

More recently, the epigenetic effects of AA have been recognized^13–16^ and implicated in its anti-tumor function^2–5,17–19^. As a specific antioxidant, AA serves as a co-factor for many Fe^2+^ and 2-oxoglutarate-dependent dioxygenases, including the methylcytosine dioxygenase TET enzymes which function to actively demethylate DNA by catalyzing the conversion of 5-methylcytosine (5-mC) to 5-hydroxymethylcytosine (5-hmC), and subsequently to 5-formylcytosine (5-fC) and 5-carboxylcytosine (5-caC). We have previously demonstrated intratumoral gain of 5-hmC in an RCC xenograft model with intravenous high-dose AA administration ^2^, and in a lymphoma syngeneic model with intraperitoneal high-dose AA administration ^3^. In the immunocompetent syngeneic model, this corresponded to increased immune cell infiltration, likely a result of demethylation-induced expression of tumor antigens ^3^. Increase in 5-hmC has also been demonstrated in peripheral blood cells with intraperitoneal high-dose AA ^5^. Aside from TET, HIF-hydroxylase mediated inhibition of HIF activity has also been shown to be involved in the anti-tumor effect of high-dose parenteral AA ^12,20,21^.

Here, we identify syngeneic models with varied sensitivity to AA, in which we determine the relative contribution of the seemingly opposing cofactor (antioxidant) activity and oxidative stress in the *in vivo* anti-tumor response to pharmacologic AA, as a single agent and in combination with anti-PD1 immunotherapy. We then shed light on factors underlying the marked difference in AA-induced effects *in vitro* and *in vivo*, with critical implications not only for the clinical translation of high-dose AA in cancer treatment, but also the field of free radical biology.

## Results

### Differential anti-tumor effects of ascorbic acid are observed across models

The anti-tumor activity of AA was compared in two immunocompetent syngeneic murine tumor models, A20 (B-cell lymphoma) and Renca (RCC). Balb/c mice were treated daily with AA (sodium ascorbate) or Vehicle (sodium chloride) beginning once subcutaneous tumors were palpable and continuing until a humane endpoint of tumor size was met. For A20 and Renca models, treatment duration was 9 and 10 days, respectively. Consistent with our previously reported study ^3^, we confirmed anti-tumor activity of single agent AA in the A20 lymphoma syngeneic model (**Fig 1A,B**). Inhibition of tumor growth was observed from the third day of AA administration and continued until tumors were harvested (P<0.05; **Fig 1A**), at which point AA treated tumors weighed 55% of the Vehicle treated group (1063 ± 224 mg vs 1936 ± 396 mg, AA vs Vehicle, P=0.05; **Fig 1B**). In contrast to the A20 model and our previous report of anti-tumor activity of AA in an RCC 786-O xenograft model ^2^, AA did not exhibit significant anti-tumor activity in the Renca model at any point in the course of treatment (P>0.05, **Fig 1C**) or in final tumor weight (824 ± 130 vs 876 ± 188, AA vs Vehicle, P=0.41; **Fig 1D**). Moreover, unlike the potentiation of anti-PD1 response by AA previously shown in the A20 model ^3^, AA did not improve anti-tumor activity of anti-PD1 in the Renca model (**Fig 1C,D**). To further investigate the potential mechanisms mediating these differential anti-tumor effects, we next sought to compare the expression of the known AA transporters, SVCT1 (encoded by*Slc23a1*) and SVCT2 (encoded by *Slc23a2*), as well as the methylcytosine dioxygensase TET enzymes (encoded by *Tet1*, *Tet2*, and *Tet3)* which we and others have previously demonstrated to mediate the epigenetic effects of ascorbic acid.

**Fig 1.**
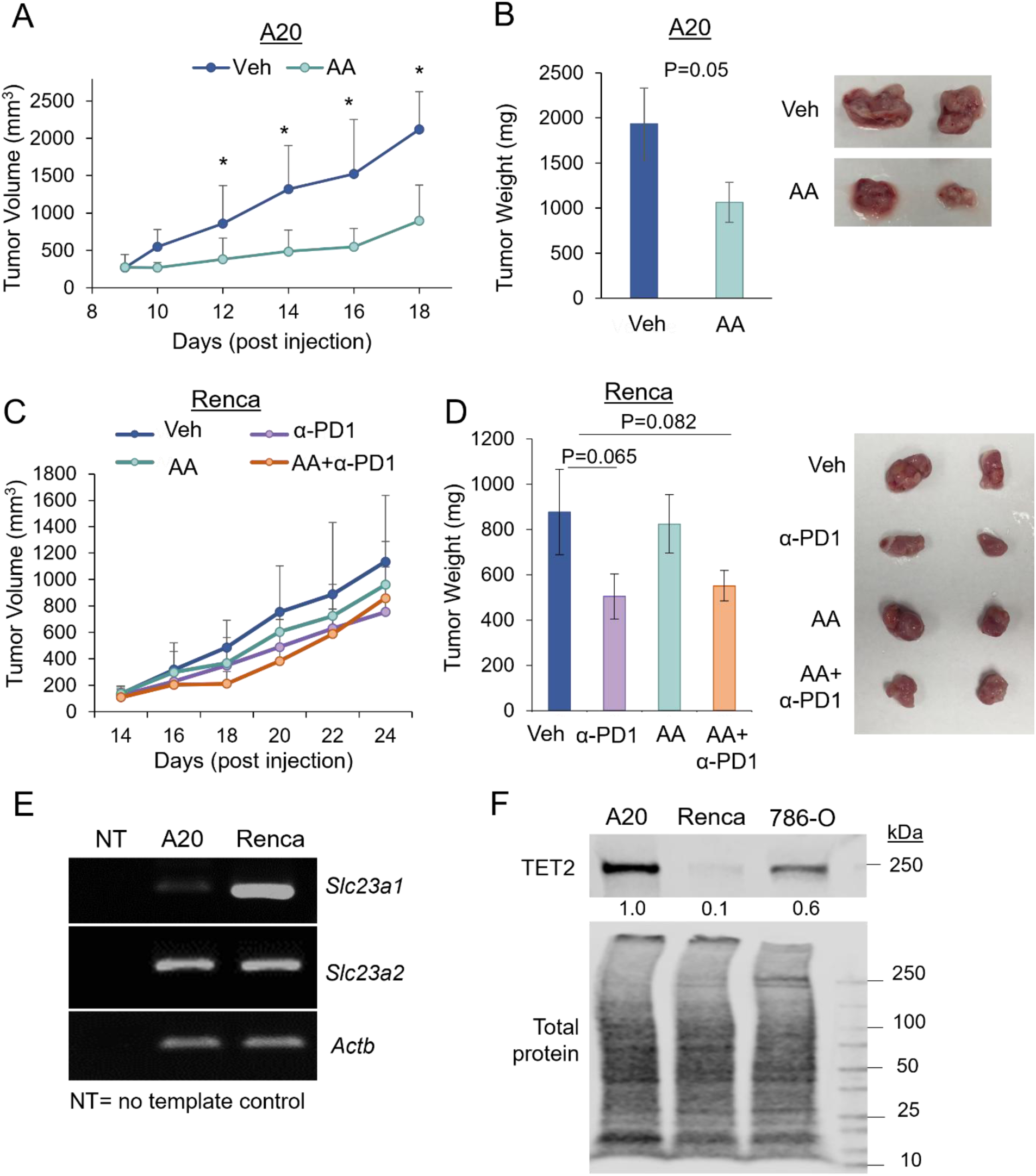
Differential anti-tumor effects of ascorbic acid are observed across syngeneic models. **A-B.** Balb/c mice bearing A20 tumors were treated daily with AA (n=6 mice) or Vehicle (Veh, n=6 mice) beginning day 9. Sensitivity to AA was observed in tumor volume plotted over the course of treatment (A; mean ± sd) and in final excised tumor weight (representative tumors shown). **C-D.** Balb/c mice bearing Renca tumors were treated with Vehicle (Veh, n=5 mice), AA (n=5 mice), anti-PD1 (n=5 mice), or AA+anti-PD1 (n=6 mice) beginning day 14. Resistance to AA was observed in tumor volume plotted over the course of treatment (C) and final excised tumor weight (representative tumors shown). **E.** PCR products for *Slc23a1*, *Slc23a2*, and *Actb* in A20 and Renca cell lines. **F.** Western blot for TET2 and SVCT1 in A20, Renca, and 786-O cell lines. Data are represented as mean ± se unless otherwise specified. P-values represent Welch’s T-test. *, p<0.05

### *Slc23a1*, *Slc23a2*, and *Tet2* are differentially expressed in A20 and Renca cell lines

Renca was found to have high expression of both *Slc23a1* and *Slc23a2* whereas A20 predominantly expressed *Slc23a2* (**Fig 1E**). Comparison of TET proteins showed that expression of TET2 was markedly lower in Renca compared to the AA-sensitive models, A20 and 786-O (**Fig 1F**). This trend was also observed for TET3, albeit to a lesser degree, while TET1 expression was detected primarily in the 786-O cell line (**Suppl Fig S1A, S1B).** Baseline 5-hmC was also lower in Renca compared to A20 cell lines, as expected with the very low TET2 expression in Renca (**Suppl Fig S1C**). We focused on *Tet2* in follow-up studies for two additional reasons: (i) in clear cell renal cell carcinoma (TCGA KIRC), expression of *TET2*, but not *TET1* or *TET3*, is negatively correlated with DNA methylation; and associated with inferior overall survival^19^; and (ii) TET2 has been shown to be a tumor suppressor in several malignancies. Similarly, we focused on *Slc23a2* in follow-up studies as it is the dominantly expressed ascorbate transporter in most tissues except kidney and gastrointestinal tract (which express *Slc23a1*), and is also the most dominantly expressed ascorbate transporter in a majority of cancers. A20 tumors too had high expression of *Slc23a2* and minimally expressed *Slc23a1* (**Fig 1E**).

### Knockdown of *Scl23A2* or *Tet2* blocks anti-tumor activity of AA in the sensitive A20 model

A20 cells with stable shRNA knockdown of *Tet2* (sh*Tet2*) and *Slc23a2* (sh*Slc23a2*) as well as non-targeting control (shNC) were generated as shown in **Fig 2A**. A dot blot for 5-hmC confirmed attenuation of 5-hmC following treatment with ascorbic acid in sh*Tet2* and to a lesser extent sh*Slc23a2* cells compared to shNC (**Fig 2B**). Cell growth was not significantly affected by *Tet2* or *Slc23a2* knockdown (**Suppl Fig S2A**). We then used these stably modified cell lines to generate syngeneic tumor grafts in Balb/c mice and treated with Vehicle or AA as in Fig 1. Similar to our previous A20 studies, AA treatment resulted in anti-tumor effect in shNC tumors (**Fig 2C,D).** Importantly, loss of either *Slc23a2* or *Tet2* was sufficient to fully block this response, with no response to AA observed throughout the treatment period (**Fig 2C,D**).

**Fig 2.**
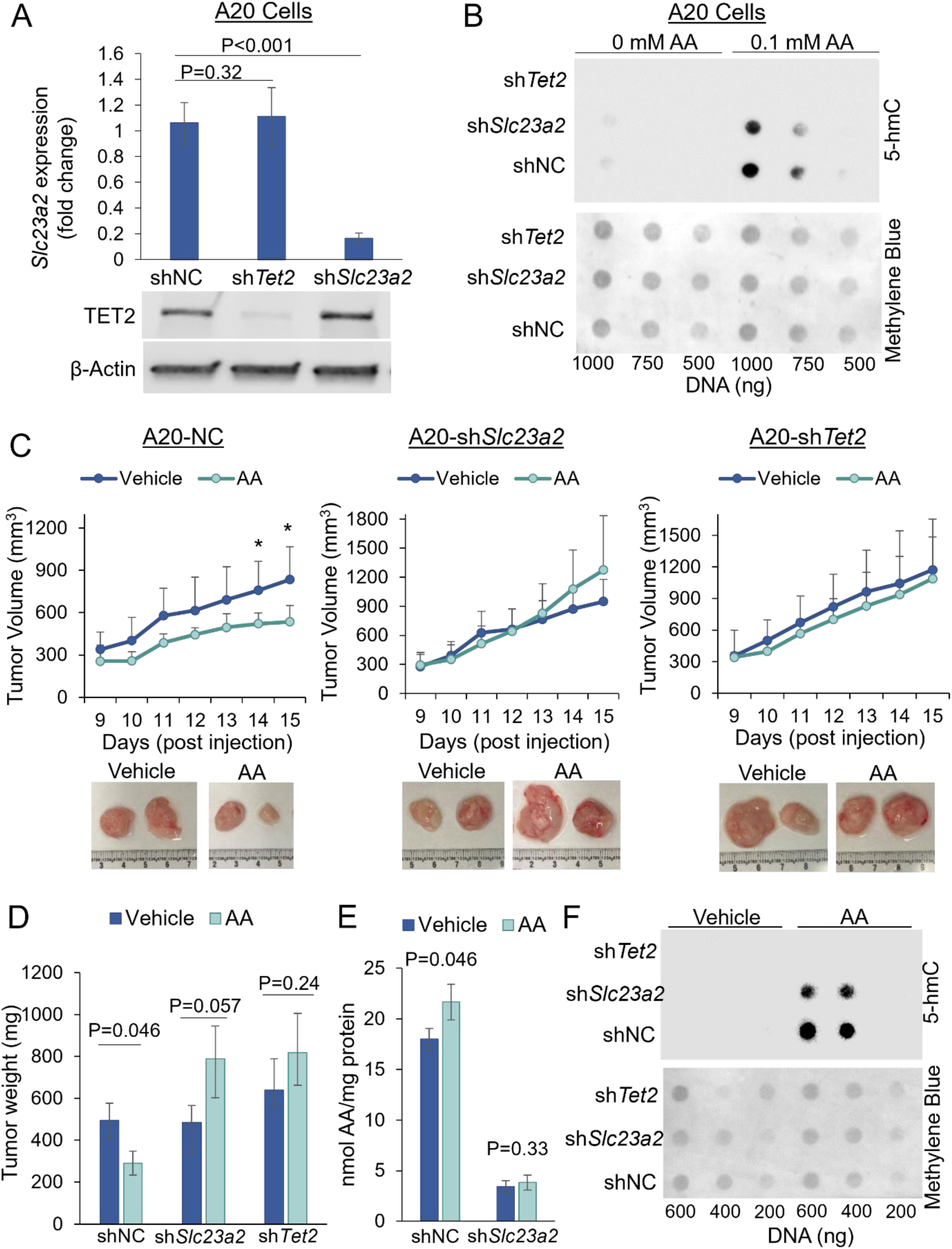
Knockdown of *Slc23a2* or *Tet2* in an AA-sensitive model ablates response to AA. **A.** PCR for *Slc23a2* and Western blot for TET2 confirmed shRNA knockdown of *Slc23a2* and *Tet2* in A20 cells. **B.** 5-hmC dot blot of A20 cell lines pre-treated with 10 µg/ml catalase for 30 min followed by 0 or 0.1 mM AA for 6 h and harvested 24 h after the start of AA treatment. Methylene blue stain is shown for loading control. **C.** Balb/c mice were injected with stably transduced A20 tumor cells as in Figure 1. Once tumors were palpable (day 9 post tumor cell injection; shNC, n=11; sh*Tet2*, n= 14; sh*Slc23a2*, n=14), mice were randomized into two treatment groups, and daily AA or Vehicle treatments began. Tumor volume was measured daily (mean ± sd). **D.** Mice were euthanized on day 16 and excised tumors were weighed. **E.** Ascorbic acid content was determined in fresh tumor lysates to compare effects of ascorbic acid treatment and *Slc23a2* transporter knockdown. Ascorbic acid was last administered 24 h prior to tumor harvest. **F.** 5-hmC dot blot of DNA isolated from A20 tumors 24 h post-Vehicle or AA administration. Representative blot shown. Data are represented as mean ± se unless otherwise specified. P-values represent Welch’s T-test. *, p<0.05

To confirm that loss of *Slc23a2* prevented ascorbate uptake, we measured intratumoral ascorbate levels. While ascorbate was elevated in AA-treated shNC tumors at the time of harvest (P=0.046; 24 h post-AA treatment), this elevation was not observed in AA-treated sh*Slc23a2* tumors (P=0.33) (**Fig 2E**). Moreover, baseline ascorbate levels were also significantly higher in shNC compared to sh*Slc23a2* (17.98 ± 1.06 v 3.42 ± 0.57 nmol AA/mg protein; **Fig 2E**), likely reflecting physiologic ascorbate derived from functional Gulo enzyme. We also assessed global DNA 5-hmC, which we have previously reported to be associated with response to AA treatment ^3^. Similar to *in vitro* AA treatment, shNC tumors treated with AA had a strong induction of 5-hmC whereas sh*Slc23a2* tumors had a more intermediate response, and sh*Tet2* tumors had no observable 5-hmC (**Fig 2F**). Together, these data confirmed that in the A20 model, both SLC23A2-mediated transport of AA as well as TET2 expression are required for high dose AA-induced anti-tumor activity and maximal AA-induced 5-hmC.

Prolyl hydroxylase and asparaginyl hydroxylase-mediated inhibition of HIF pathway/transcriptional activity are functions of AA co-factor activity with potential anti-tumor effects. Hydroxylation of specific proline residues by HIF PHD targets HIF-α for proteasomal degradation, whereas hydroxylation of an asparagine in the C-terminal transactivation domain by asparaginyl hydroxylase (FIH) prevents its interaction with transcriptional coactivator p300. To assess whether AA was also acting on this pathway in the sensitive A20 model, we measured HIF target genes, *Vegfa*, *Glut1 (*encoded by *Slc2a1)*, and *Car9*, in shNC and sh*Slc23a2* tumors (**Suppl Fig S3A**). In the shNC group, expression of all HIF target genes was downregulated in AA-treated mice compared to Vehicle. Knockdown of *Slc23a2* in the sh*Slc23a2* group blocked this effect of AA (**Suppl Fig S3B**). Thus, downregulation of the HIF pathway may be an additional co-factor function of AA contributing to its anti-tumor activity in the A20 model.

### *Tet2* reconstitution inhibits tumor growth and enhances AA sensitivity in the resistant Renca model

Given the high baseline *Slc23a1* and *Slc23a2* expression and low TET2 expression in Renca cells, we next sought to assess whether reconstitution of *Tet2* would result in enhanced sensitivity to AA. For this, stable *Tet2* or empty vector-expressing Renca clones were generated (**Fig 3A**). As expected, overexpression of *Tet2* resulted in greater 5-hmC abundance following AA treatment (**Fig 3B**). These Renca clones were then used for syngeneic tumor graft induction and AA treatment as in Fig 1. *Tet2* overexpression led to significant inhibition of tumor growth *in vivo*, irrespective of AA treatment (**Fig 3C,D**). AA-responsiveness was observed in the *Tet2*-expressing tumors early in the course of treatment (day 3), however tumor growth regressed over time even without AA treatment, resulting in no significant AA effect at the time of harvest (**Figs 3C,D**). Of note, *Tet2* overexpression had a much lesser effect on Renca cell growth *in vitro* (**Suppl Fig S2B**), indicating a role for immune cell infiltration in the presence of TET2, as we have previously shown in the A20 model. Consistent with high expression of *Slc23a1* and *Slc23a2* in Renca, intratumoral ascorbate levels were significantly elevated following ascorbate treatment at the time of harvest (P<0.001; 2.5 h post AA treatment) (**Fig 3E**). Unlike the A20 model when tumors were harvested 24 h after the last AA treatment, the Renca model tumors were harvested ∼2.5 h after AA treatment. Tumor 5-hmC was induced strongly in *Tet2* overexpressing tumors, irrespective of AA administration. In Empty vector tumors, AA treatment resulted in very minor 5-hmC induction (**Fig 3F**). (Unlike *in vitro*, the 5-hmC induction with TET2 overexpression even without AA treatment was strong intratumorally *in vivo*, likely due to physiologic ascorbate levels with the functional Gulo enzyme.) We also measured HIF pathway genes in the Renca Empty vector tumors as a secondary measure of AA co-factor activity (**Suppl Fig S3C**). Expression of HIF target genes, *Vegfa*, *Glut1*, and *Car9* was not downregulated by AA treatment. To test the role of TET2 in CD8 T cell infiltration, we assessed CD8 positivity by IHC in the Renca resistant model with and without *Tet2* overexpression. Overexpression of *Tet2* increased CD8 T cell infiltration (% of total cells; Empty vs Tet2, 0.63 ± 0.20 % vs 1.48 ± 0.37 %, p=0.031) (**Fig 3G, Suppl Fig S4**).

**Fig 3.**
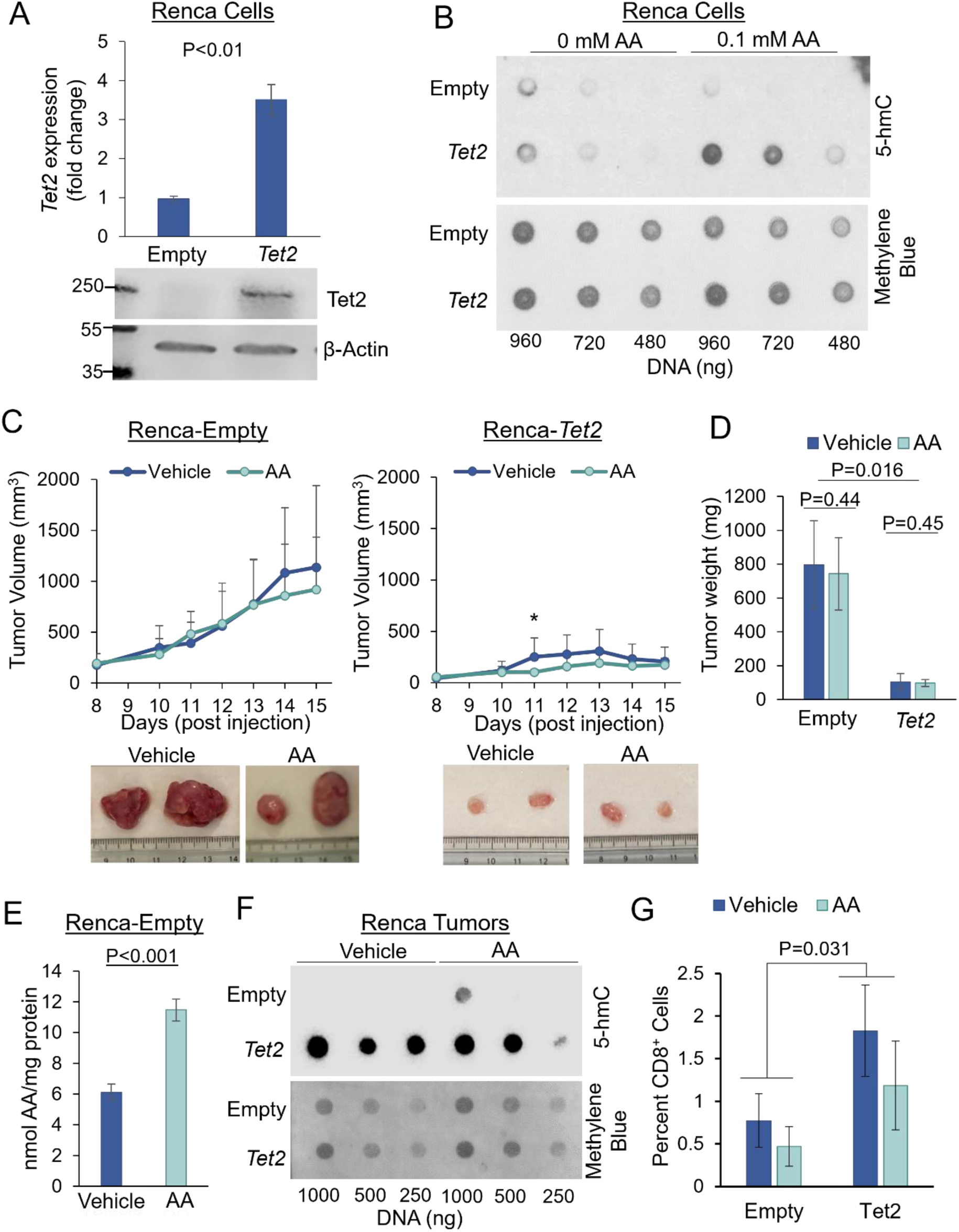
Overexpression of Tet2 in an AA-resistant model markedly attenuates tumor growth and enhances response to AA. **A.** PCR and Western blot confirmed overexpression of *Tet2* in Renca cells. **B.** 5-hmC dot blot of Renca modified cell lines pre-treated with 10 µg/ml catalase for 30 min followed by 0 or 0.1 mM AA for 6 h and harvested 24 h after the start of AA treatment. Methylene blue stain is shown for loading control. **C.** Balb/c mice were injected with stably transduced Renca tumor cells as in Figure 1. Once tumors were palpable (day 8 post tumor cell injection; Empty, n= 16; *Tet2*, n= 13), mice were randomized to AA or Vehicle treatments and daily treatments began. Tumor volume was measured daily (mean ± sd). **D.** Mice were euthanized on day 16 and excised tumors were weighed. **E.** Ascorbic acid content was determined in fresh tumor lysates to compare effects of ascorbic acid treatment. Ascorbic acid was administered 2.5 h prior to tumor harvest. **F.** 5-hmC dot blot of Renca tumors 2.5 h post Vehicle or AA administration. Representative blot shown. **G.** Immunohistochemistry for CD8 in Renca tumor model with and without *Tet2* overexpression shows increased infiltration of CD8^+^ cells with *Tet2* overexpression (p=0.031). Representative IHC images shown in Suppl Fig S4. Data are represented as mean ± se unless otherwise specified. P-values represent Welch’s T-test. *, p<0.05

### High-dose AA treatment *in vivo* resulted in increased intratumoral antioxidant capacity without evidence of oxidative stress in both sensitive A20 and resistant Renca models

Using the A20 and Renca tumor models of varied sensitivity to AA, we further explored whether responsiveness to AA was associated with antioxidant activity and/or oxidative stress. We measured the total antioxidant capacity of fresh tumor samples, quantified by reduction of Fe^3+^ to Fe^2+^. In AA-sensitive A20 shNC tumors, AA treatment significantly increased intratumoral antioxidant capacity (**Fig 4A**). In contrast, antioxidant capacity was not affected by AA treatment in AA-insensitive sh*Slc23a2* tumors (**Fig 4A**). To determine whether catalase expression was influenced by AA treatment and therefore contributing to antioxidant function, we assessed catalase protein expression in tumors (**Fig 4B**). Catalase expression was not affected by AA treatment or knockdown of *Slc23a2* or *Tet2*. Together, these data provide strong support for parenteral AA increasing *in vivo* intratumoral antioxidant capacity.

**Fig 4.**
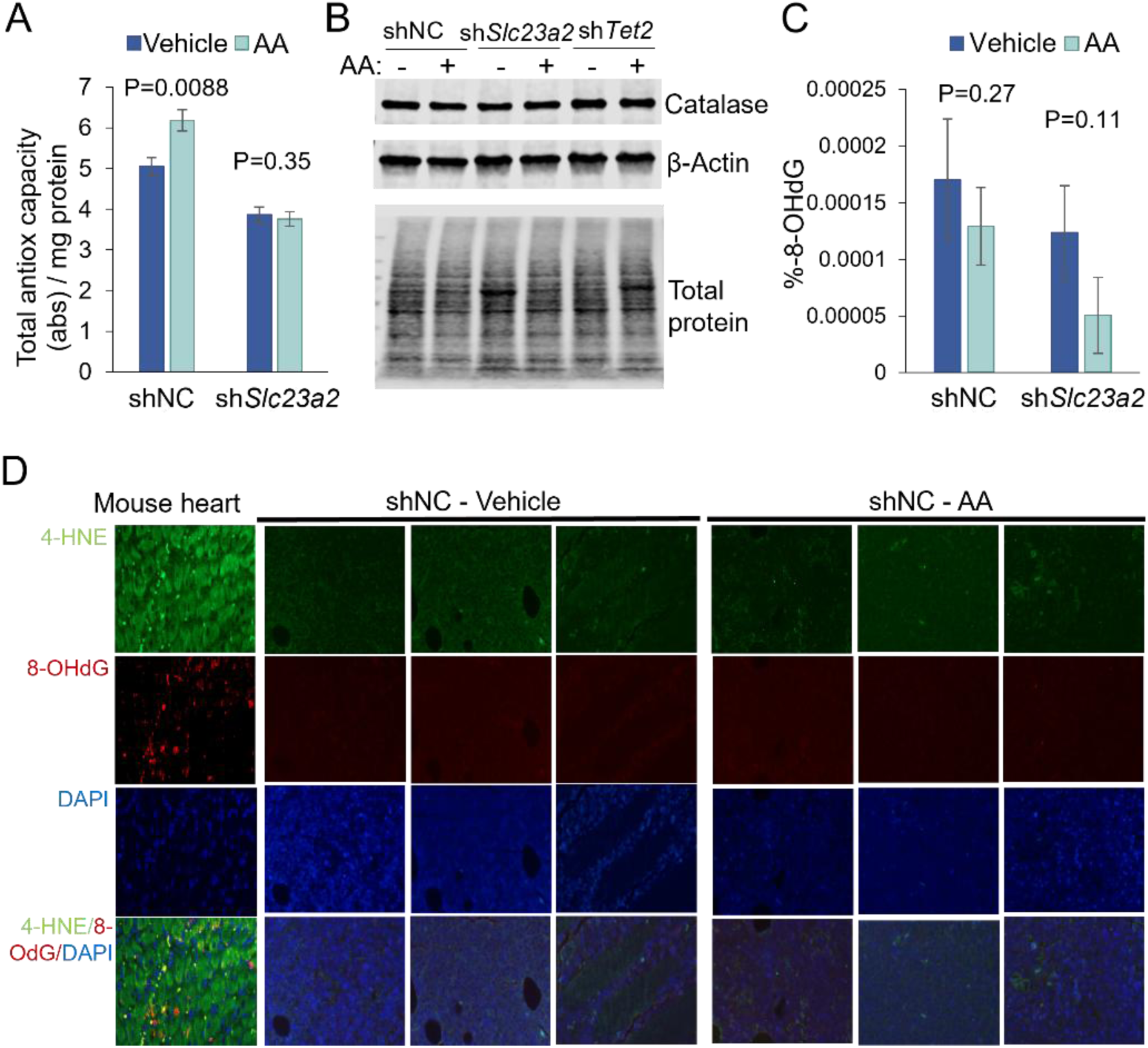
In an AA-sensitive model, high-dose AA increases intra-tumoral antioxidant capacity and does not induce oxidative stress. **A.** Total antioxidant capacity was measured in fresh tumor tissues. **B.** Catalase expression is unchanged by ascorbic acid treatment and knockdown of *Slc23a2* or *Tet2*. Representative blot shown. **C.** 8-OHdG % was measured in DNA extracted from flash-frozen tumors and determined by ELISA (n=4 per group). Data represented as mean ± se, P-values represent 1-sided Welch’s T-test. **D.** Immunofluorescence for 8-OHdG and 4-HNE was performed in FFPE tumors from shNC mice treated with Vehicle and AA show negativity for both oxidative marks. Tumors with median tumor size per group (n=3) are shown. Mouse heart was used as a positive control.

We next sought to assess whether high-dose AA also led to a pro-oxidant effect in this model using two well-established markers of oxidative stress: the DNA oxidative damage marker, 8-OHdG, and the lipid peroxidation marker, 4-HNE. An 8-OHdG ELISA using tumor DNA showed that tumors were largely negative for the 8-OHdG mark with no significant effect of AA treatment (**Fig 4C**). Tissue immunofluorescence for 8-OHdG and 4-HNE confirmed negativity for 8-OHdG and very low baseline 4-HNE positivity across all shNC and sh*Slc23a2* tumor samples (**Fig 4D**, **Suppl Fig S5**).

We also characterized antioxidant vs pro-oxidant function of AA in the Renca model, in which tumors were harvested 2.5 hours after AA administration, unlike the A20 model where tumors were harvested 24 hours later. Similar to A20, AA treatment resulted in a significant increase in antioxidant activity in Renca (**Fig 5A**). Catalase expression was consistent across tumors and was unchanged by AA treatment (**Fig 5B**). Also similar to the A20 model, 8-OHdG was not affected by AA treatment (**Fig 5C**). Tissue immunofluorescence confirmed lack of 8-OHdG and 4-HNE in Renca tumors (**Fig 5D**). Taken together, both A20 and Renca tumor models consistently demonstrated an increase in intratumoral antioxidant capacity without evidence of oxidative stress with parenteral high-dose AA treatment.

**Fig 5.**
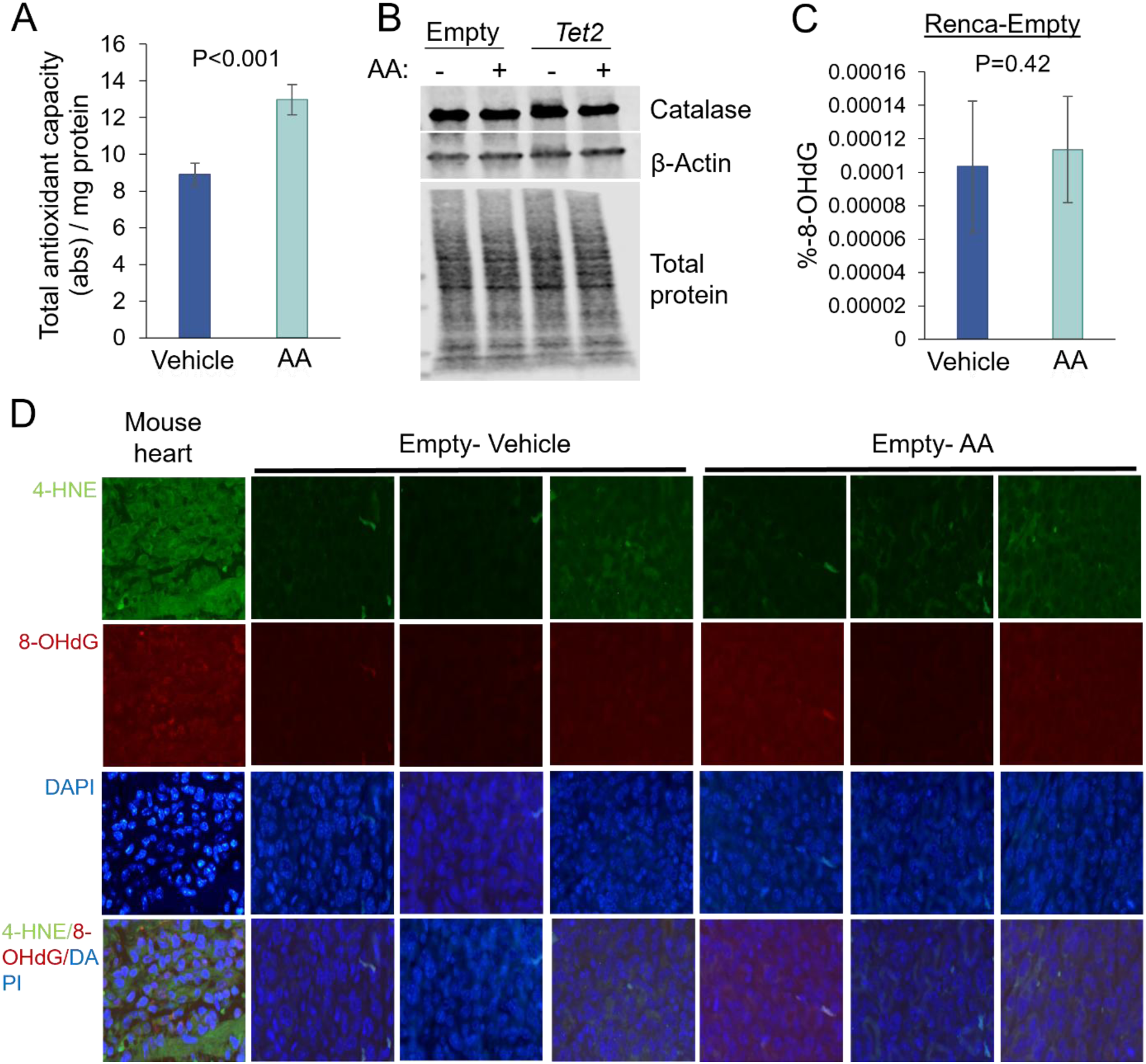
In an AA-resistant model too, high-dose AA increases intra-tumoral antioxidant capacity and does not induce oxidative stress. **A.** Total antioxidant capacity was measured in fresh tumor tissues. The assay was performed on all tumors. **B.** Tumor Catalase expression is unchanged by ascorbic acid treatment or overexpression of *Tet2*. Representative blot shown. **C.** 8-OHdG % was measured in DNA extracted from flash-frozen tumors and determined by ELISA (n=4 per group). Data represented as mean ± se, P-values represent 1-sided Welch’s T-test. **D.** Immunofluorescence for 8-OHdG and 4-HNE was performed in FFPE tumors from Empty vector mice treated with Vehicle and AA show negativity for both oxidative marks. Tumors with median tumor size per group (n=3) are shown. Mouse heart was used as a positive control.

Of note, in both these models, parenteral high-dose AA administration achieved millimolar intratumoral AA concentrations (10nmol AA/mg protein amounts to 2mM AA with an estimate of 200g/L intracellular protein concentration).

### Unlike intra-tumoral AA function *in vivo,* high-dose AA treatment *in vitro* resulted in robust oxidative damage in A20 and Renca cells

Oxidative stress has been proposed as an anti-tumor function of AA based on a large number of *in vitro* findings. To confirm the differential activity of AA *in vivo* vs *in vitro,* we performed AA treatment *in vitro* for A20 and Renca cell lines and measured oxidative stress. In line with *in vitro* literature, treatment with high-dose AA (1 mM) in both A20 and Renca cell lines resulted in significant induction of oxidative stress, as evidenced by positive 8-OHdG and 4-HNE staining (**Fig 6A,B**). Treatment with 1 mM H_2_O_2_ served as a positive control. Pre-treatment of cells with catalase blocked the pro-oxidant function of AA (**Fig 6A,B**) as shown previously, again demonstrating that H_2_O_2_ is the mediator of AA-induced oxidative stress *in vitro*. (AA millimolar concentrations that did not result in any evidence of oxidative damage intratumorally *in vivo,* caused robust oxidative damage *in vitro.*)

**Fig 6.**
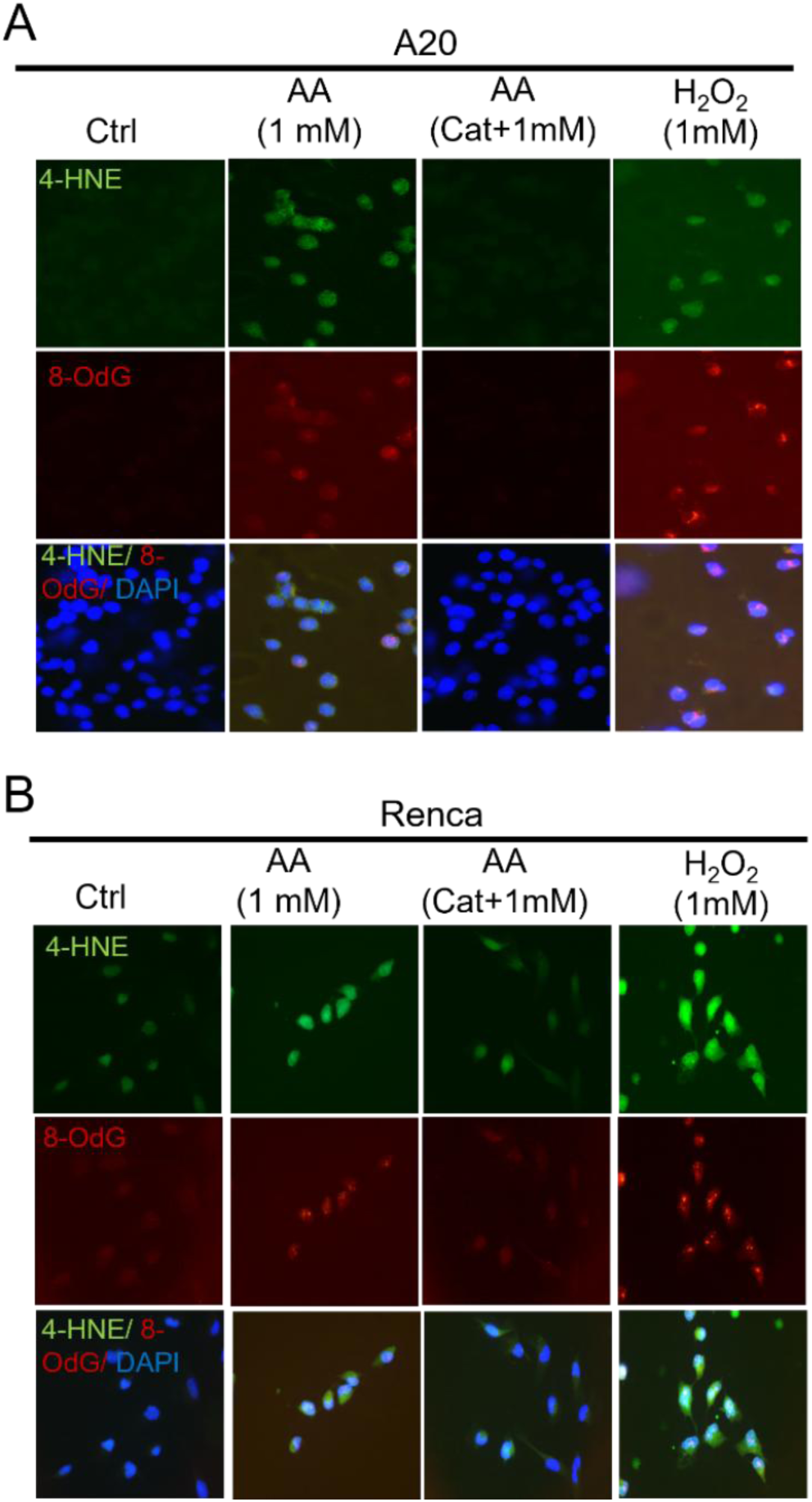
Unlike intra-tumoral AA function *in vivo*, high-dose AA induces robust oxidative stress *in vitro*. **A-B.** Cells were treated with 1 mM ascorbic acid or 1 mM H_2_O_2_ for 4 h (A20) or 6 h (Renca). Catalase (10 µg/ml) was added 30 minutes prior to ascorbic acid treatment as a negative control. Oxidative markers 8-OHdG and 4-HNE were assessed by immunofluorescence.

### AA-induced H_2_O_2_ production is markedly influenced by molecular oxygen concentration, underlying the discrepancy between abundant *in vitro* and absent *in vivo* oxidative effects of high-dose AA

We then investigated the underlying mechanisms for the discrepancy between the abundant *in vitro* and absent *in vivo* oxidative stress with high-dose AA.

The following reactions are involved in the production of H_2_O_2_ with AA *in vitro*.

1. Ascorbate reduces catalytic metal ions such as ferric ions to ferrous ions.

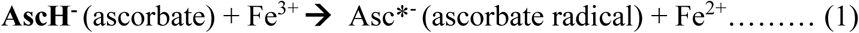
2. The ferrous ions react with oxygen to form superoxide radical

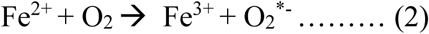
3. Superoxide radicals then dismutes to H_2_O_2_ and O_2_

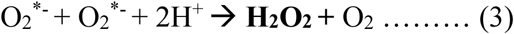

Hydrogen peroxide can spontaneously decompose to water and oxygen (and the reaction is catalyzed by the enzyme catalase)

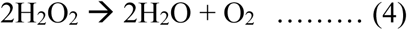

We hypothesized that reaction 2 involving molecular oxygen may be the reason for the discrepancy (molecular oxygen in regular *in vitro* cell culture conditions is present at supraphysiologic levels of ⁓19%, whereas mammalian tissue oxygen concentration is much lesser at 1-5%). The ascorbate radical produced in reaction 1 is relatively less reactive, undergoing an equilibrium reaction to form dehydroascorbic acid and ascorbate: 2Asc*^-^ + H+ ←→ DHA + AscH-. We reasoned that the reactive ions (such as Fe^2+^) generated from reaction 1 could massively trigger reaction 2 with the high availability of molecular O_2_ *in vitro*, resulting in an abundance of superoxide ions and subsequent generation of H_2_O_2._ The decomposition of H_2_O_2_ to water and oxygen (reaction 4) may also be reduced in the presence of supraphysiologic oxygen concentration.

To test the hypothesis that difference in molecular O_2_ concentration is the main reason for the discrepancy, we exposed cell lines to high-dose AA in both regular (19%) and hypoxia (3%) incubators, the latter representing *in vivo* physiologic O_2_ concentration, with ***dynamic real-time*** measurements of H_2_O_2_ at 30 min intervals over 10 hours **(Fig7)**. With high-dose AA treatment of Renca cells (1mM and 5mM), both the rate of H_2_O_2_ production and the total amount of H_2_O_2_ produced was markedly higher under normoxia conditions compared to hypoxia **(Fig 7A, 7B)**. H_2_O_2_ (0.1mM) served as a positive control for this experiment, which spiked immediately upon administration and did not increase over time **(Fig 7C)**. Of note, the H_2_O_2_ signals with 1mM AA under hypoxic conditions were far lesser than that with 100µM H_2_O_2_. (Interestingly, even with the 0.1mM H*_2_*O_2_ control, the measured H_2_O_2_ was lower in hypoxia likely due to reaction 4 above i.e. more spontaneous decomposition to water and oxygen.) In line with these results, cell viability measured by Cell Titer Blue following short-term high-dose AA treatment was also dependent on molecular O_2_ concentration. In Renca cells, treatment with 5 mM AA for 4 hr under normoxic conditions resulted in 13% viability while the same treatment under hypoxic conditions resulted in 62% viability **(Fig 7D)**. Likewise, treatment of A20 cells with 5 mM AA for 2 hr under normoxic compared to hypoxic conditions yielded 2.8% vs 69% viability **(Fig 7D)**. Importantly, reduction in cell viability was fully reversed in the presence of catalase in both conditions, confirming H_2_O_2_-mediation **(Fig 7D-E)**.

**Fig 7.**
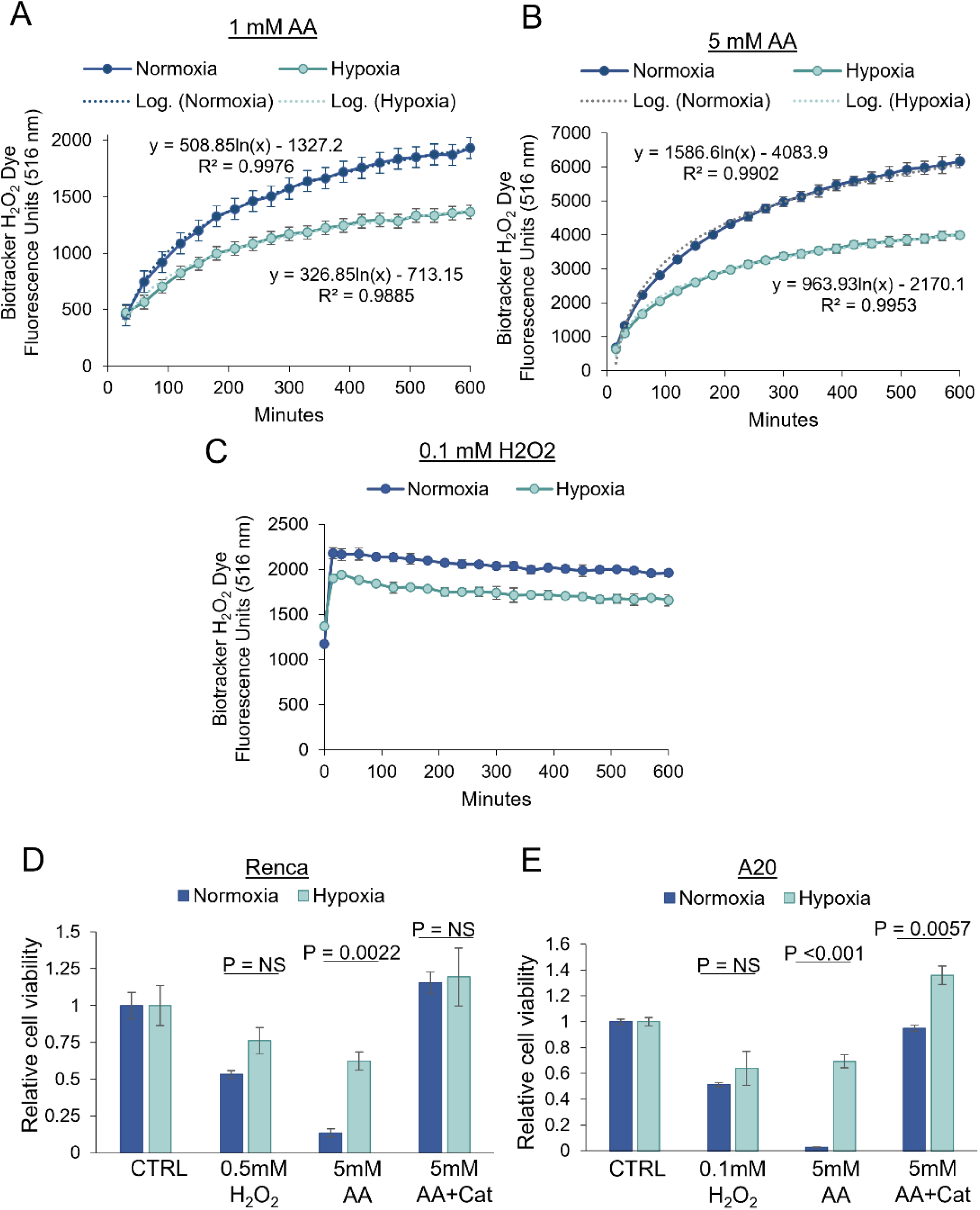
AA-induced H_2_O_2_ production is markedly influenced by molecular oxygen concentration, underlying the discrepancy between abundant *in vitro* and absent *in vivo* oxidative effects of AA. **A-B.** H_2_O_2_ dye was added to Renca cells plated in a 96-well plate, and cells were allowed to equilibrate to normoxic (19% O_2_) or hypoxic (3%) conditions. Cells were then treated with 1 mM or 5 mM AA or 0.1 mM H_2_O_2_ under atmospheric conditions and immediately returned to their respective normoxic or hypoxic conditions for the remainder of the experiment. H_2_O_2_ was measured every 15 to 30 minutes over the course of a 10-hour period. Data are plotted as mean ± sd for each timepoint. Logarithmic curves were fit to the AA-induced H_2_O_2_ generation over time, in which the maximum production of H_2_O_2_ (1 mM: 1933.00 ± 50.27 vs 1364.70 ± 24.21; 5 mM: 6161.67 ± 199.76 vs 3987.67 ± 112.88 fluorescence units) were much higher under normoxic compared to hypoxic conditions. **C**. H_2_O_2_-treated cells served as a positive control for H_2_O_2_ measurement. **D-E.** Renca and A20 cells were treated with AA, H_2_O_2_, and/or catalase (Cat) under normoxia (19% O_2_) or hypoxia (3% O_2_) conditions. Cell viability was determined after 4 h (Renca) or 2 h (A20) of treatment. Data are represented as means ± se. Statistical comparisons performed by Student’s T-test.

Apart from the marked difference in molecular oxygen concentration and subsequent H_2_O_2_ generation with high-dose AA, it is possible that the difference in cell density in tumors and tumoral architecture *in vivo* versus cells *in vitro* confer a markedly different overall ability to counter H_2_O_2_-induced oxidative effects. We used Renca cells which grow in clusters to study the impact of cell density and confluence pattern on H_2_O_2_-induced oxidative stress and cytotoxicity (**Suppl Fig S6, S7**). With lower pre-treatment cell density, cell viability was markedly reduced and oxidative damage markers 8-OHdG and 4-HNE were induced to a much greater extent, compared to higher pretreatment cell density, with the same H_2_O_2_ exposure (1mM over 2hr) **(Suppl Fig S6A,B)**. With regards to growth pattern, there was marked reduction in viability and increase in oxidative stress with scattered pattern compared to clustered pattern of growth in Renca, despite similar cell count at treatment (0h) and similar viability of respective controls **(Suppl Fig S7A-D,** experimental design in methods). Together, these experiments indicate an increased ability of tumors *in vivo* (with higher cell density and a dense/clustered architecture) to neutralize *any* small amounts of extracellular H_2_O_2_ compared to cells *in vitro*.

Finally, as a positive control for *in vivo* oxidative stress, we sought to measure intra-tumoral oxidative stress following doxorubicin treatment. Oxidative stress is one of the well described consequences of doxorubicin treatment and has been extensively shown both *in vitro* as well as *in vivo* ^22–24^. For this study, we treated A20 tumor-bearing mice with vehicle (Veh) or doxorubicin (Dox) beginning once tumors were palpable. Tumor volume and final tumor weight showed an anti-tumor effect (**Fig 8A,B**). To assess oxidative stress, we used markers for both lipid (MDA) and DNA oxidative damage (8-OHdG) that have been previously shown to be increased by Dox *in vitro* and in heart tissue^23–25^. Indeed, both MDA and 8-OHdG were elevated in tumors of Dox-treated mice (**Fig 8C,D**), unlike with high-dose parenteral AA. We then performed a follow-up study treating A20 bearing mice with Vehicle, Dox, or the combination of Dox and high-dose AA to test whether the addition of AA would affect the anti-tumor activity of Dox via antioxidant function. The addition of high-dose AA to Dox did not affect tumor size compared to Dox treatment alone (**Fig 8E,F**). This suggests that the anti-tumor effect of Dox is mediated primarily via non-oxidative mechanisms. Moreover, mean 8-OHdG% in tumors was reduced with the addition of AA (**Fig 8G**, Veh, 0.0058% ± 0.00057%; Dox, 0.0076% ± 0.00072%; AA+Dox, 0.0057% ± 0.00082%).

**Fig 8.**
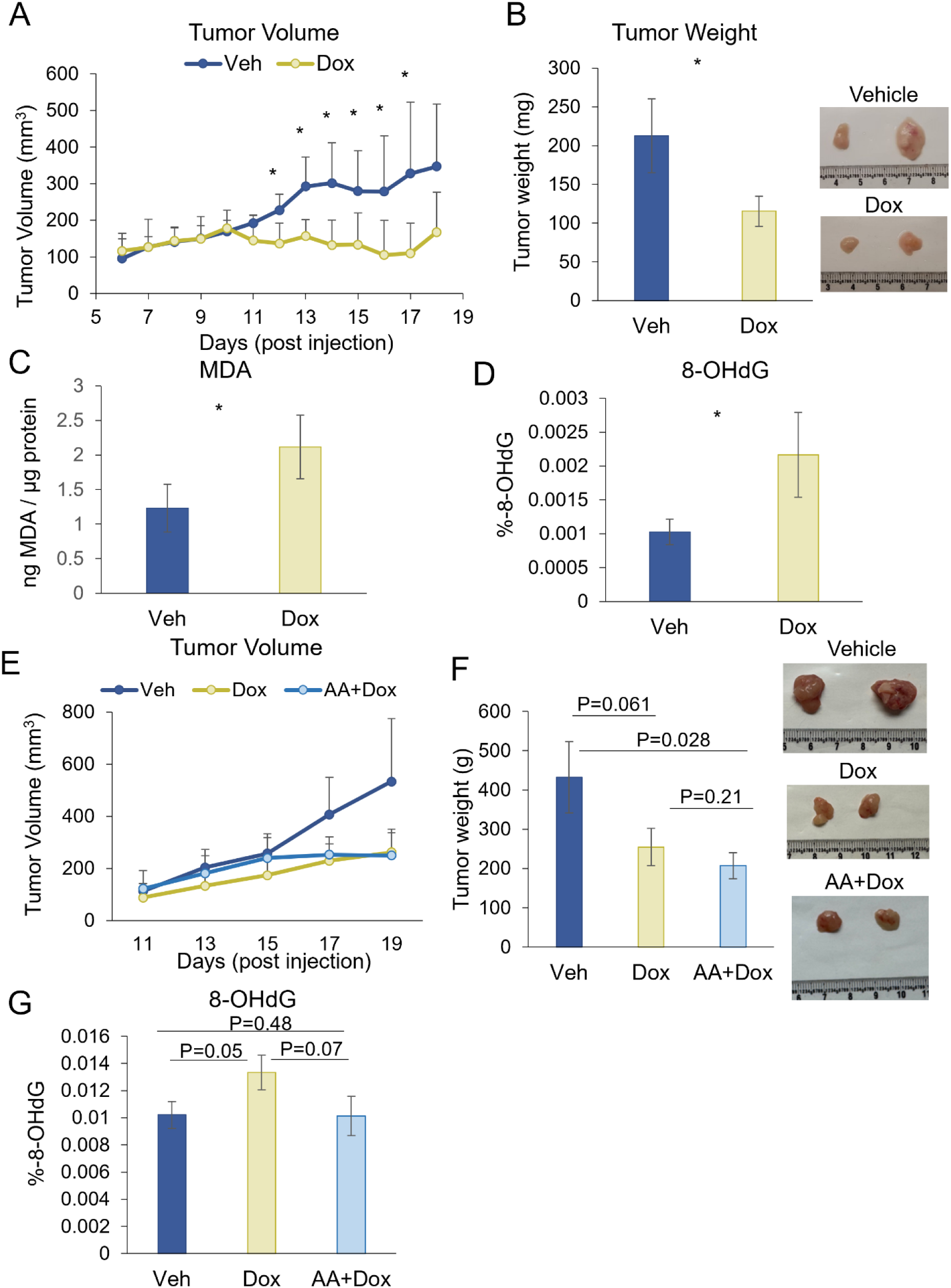
Doxorubicin treatment results in oxidative stress *in vivo*. **A-D.** Balb/c mice bearing A20 tumors were injected with 1 mg/kg doxorubicin (Dox) or vehicle (Veh) once tumors were palpable (n=4 mice per treatment). Treatments continued every 3 days until animals reached a humane endpoint, 19 days post-tumor cell injection. **A.** Tumor volume was measured daily (mean ± sd). **B.** Tumor weight was measured at tumor harvest, 19 days post-tumor cell injection (P=0.05). **C.** MDA lipid peroxidation analysis and **(D.)** 8-OHdG DNA damage analysis was performed on all mouse tumors (P= 0.087 and P=0.066, respectively). *, p<0.1. **E-G.** Balb/c mice with A20 tumors were treated with Veh (n=6), Dox (n=9), or AA+Dox (n=9). **E.** Tumor volume was measured every 2 days (mean ± sd). **F.** Endpoint tumor weight, p-values represent T-test. **G.** Tumor 8-OH-dG% was determined on n=4 per treatment, p-values represent Welch’s T-test. Data represent mean ± se unless otherwise specified.

### Low catalase does not confer sensitivity to high-dose AA *in vivo*, and TET2 is not sufficient for AA-induced anti-tumor activity or potentiation of anti-PD1

We have previously shown in the sensitive A20 model that high-dose AA increased CD8 T cell infiltration, leading to potentiation of anti-PD1 therapy^3^, and we have now demonstrated the abrogation of AA’s anti-tumor efficacy in this model with stable *Tet2* knockdown. We have also demonstrated that in the Renca model, with very low/absent TET2 expression, high-dose AA is not effective as a single agent and does not potentiate anti-PD1, with *Tet2* overexpression leading to significant inhibition of tumor growth *in vivo*, irrespective of AA treatment. We have demonstrated the absence of intra-tumoral oxidative stress with high-dose AA in both models.

To further investigate the role of TET2 and *any* potential H_2_O_2_-induced oxidative stress, we next assessed the role of high-dose AA, alone and in combination with anti-PD1, in the MB49 model. MB49 has similar TET2 expression to A20 but much higher than Renca (**Fig 9A**). However, unlike Renca, MB49 has very low catalase expression (**Fig 9A**), making it a particularly ideal model to test anti-tumor activity of AA by *any* potential H_2_O_2_-induced oxidative damage. Despite similar TET2 expression, baseline 5-hmC abundance is mildly lower in MB49 compared to A20 (**Fig 9B**). *Tet2* variants were not identified in MB49 to explain the mildly lower baseline 5-hmC.

**Fig 9.**
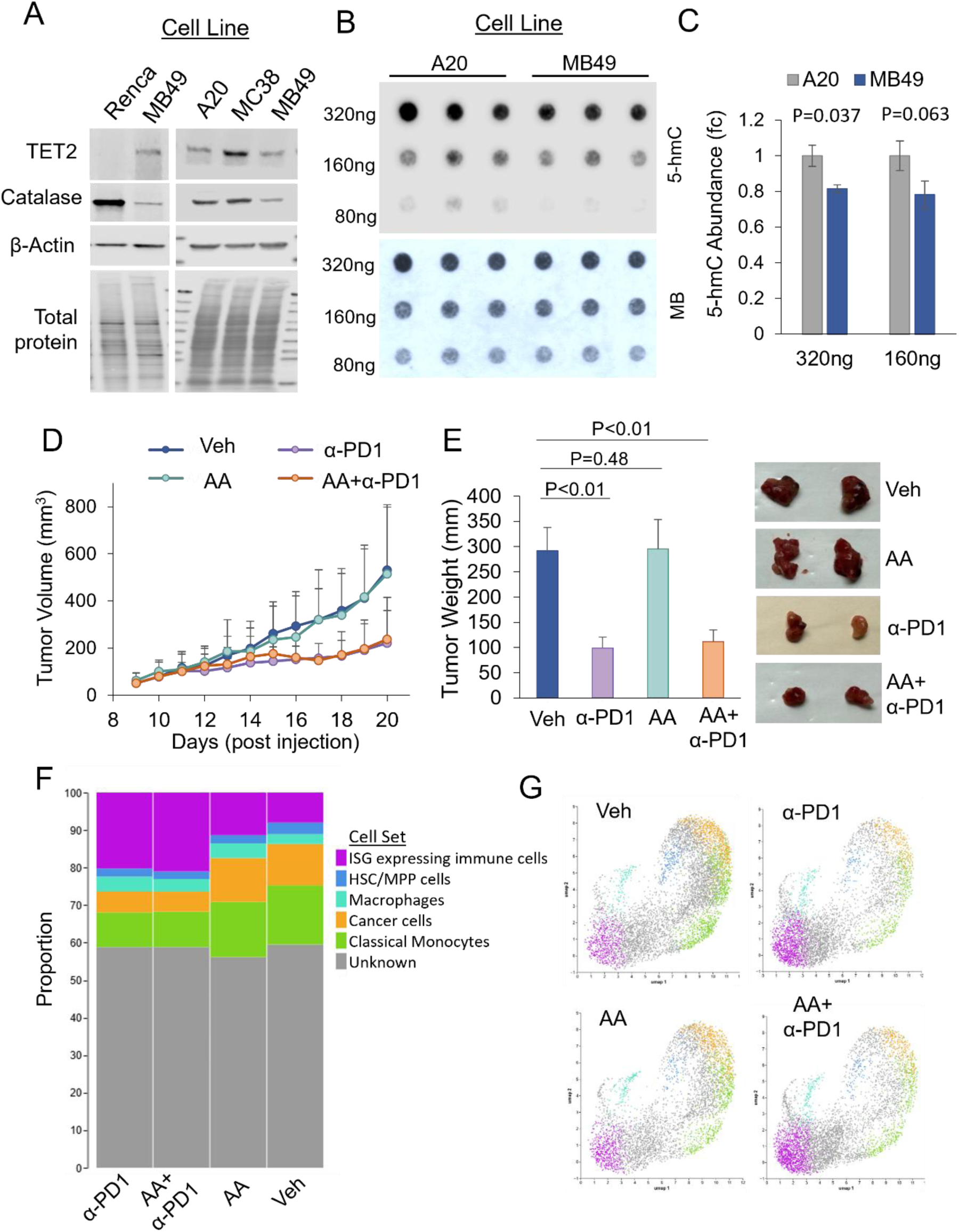
The MB49 tumor model with moderate TET2 and low Catalase expression is resistant to high-dose AA. **A.** TET2 and catalase protein expression was compared between A20, Renca, MB49, and MC38 cell lines by Western blot. **B.** Baseline 5-hmC abundance was compared between A20 and MB49 cell lines. **C.** Quantification of 5-hmC (mean ± se). **D.** Balb/c mice bearing MB49 tumors were treated daily with Vehicle (n=9), high-dose AA (n=8), anti-PD1 (n=10), or AA+anti-PD1 (n=10). Tumor volume was measured daily (mean ± sd). **E.** Tumor weight was measured at tumor harvest (mean ± se). Representative tumors shown. **F.** Single nuclei analysis shows proportion of immune cell types in MB49 tumors and (**G**) umap categorical embedding of clusters (n=4 representative tumors per treatment). P-values represent Welch’s T-test.

MB49 syngeneic tumors were established in C57/Bl6 mice to test combined treatment of AA and anti-PD1 in this model. Once tumors were palpable, mice were treated with Vehicle, AA, anti-PD1, or AA+anti-PD1. Treatment of MB49 with high-dose AA single agent did not result in tumor growth inhibition at any timepoint (**Fig 9D,E**). MB49 tumors are moderately sensitive to anti-PD1; however, AA did not potentiate anti-PD1 anti-tumor activity (**Fig 9D,E**). Single nuclei RNA-seq analysis of MB49 tumors also showed a robust anti-PD1 immune response characterized by a larger interferon-stimulated genes (ISG) expressing immune cell population (**Fig 9F,G**). No effects of AA were observed on immune cell populations in this model (**Fig 9F,G**). If H_2_O_2_ was an important anti-tumor mechanism of high-dose AA *in vivo*, MB49, with its very low catalase expression at baseline, it would be expected to be exquisitely sensitive to high-dose AA, instead of being fully resistant to single agent activity and potentiation of anti-PD1. This complete lack of response in the MB49 model despite its very low catalase expression provided further compelling evidence against the oxidative stress mechanism. However, the MB49 model also showed that TET2 expression alone is not sufficient either for single agent activity of high-dose AA or for potentiation of anti-PD1 efficacy.

We next assessed the high dose AA and anti-PD1 combination in the MC38 syngeneic model, which has very high TET2 expression and moderate catalase expression (**Fig 9A**). While MC38 exhibited significant anti-tumor activity in AA+anti-PD1 treated mice, there was no/marginal responsiveness to single agent AA (**Suppl Fig S8A-C).** Because the observed effect of combined AA and anti-PD1 therapies was greater than the expected additive effect (as seen in **Suppl Fig S8A,B**) we applied the coefficient of drug interaction (CDI) formula to calculate synergy using mean tumor volume measurements. The CDI between high-dose AA and anti-PD1 using the final mean tumor volume measurements was 0.65, indicating a significant synergistic effect (CDI < 0.7), similar to that demonstrated with the combination in the A20 model. The MC38 model revealed that even in the absence of significant single agent activity, AA can markedly potentiate anti-PD1 efficacy.

## Discussion

This study was undertaken to address the two most pressing questions in the field of ascorbic acid in cancer treatment: (i) Does high-dose parenteral AA result in *concurrent intratumoral* pro-oxidant and antioxidant/co-factor/epigenetic activity in both sensitive and resistant models? (ii) Are the biologic effects seen with high-dose AA treatment in cancer cells *in vitro* also seen in intratumoral cancer cells *in vivo*? If not, what are the mechanisms underlying the discrepancy?

### Specific antioxidant/cofactor activity with high-dose parenteral ascorbic acid *in vivo*

We show that the anti-tumor function of AA is dependent on both *Tet2* and *Slc23a2* expression in an AA-sensitive syngeneic B-cell lymphoma model, indicating critical roles for both specific antioxidant/co-factor activity and cellular AA transport. These findings are supported in the AA-resistant Renca (syngeneic renal cancer) model with high baseline AA transporter expression and absent TET2 expression, in which we reported an anti-tumor effect upon reconstitution of *Tet2*. We have previously shown that AA treatment results in increased tumor 5-hmC and recruitment of immune cells to the tumor microenvironment in the A20 model ^3^. This immune cell infiltration is potentially a result of AA-induced increase in tumor antigen expression such as human endogenous retroviruses ^3^, among other possible mechanisms. In this study, the over-expression of *Tet2* in Renca cells resulted in mild growth inhibition *in vitro* compared to the dramatic growth inhibition *in vivo. Tet2* overexpression was accompanied by increased infiltration of CD8^+^ cells, providing further evidence for the likely role of immune recognition in the TET2-dependent anti-tumor response to AA. However, our data with the MB49 (syngeneic bladder cancer) model demonstrates that TET2 expression alone is not sufficient to drive an AA-induced anti-tumor response - either as a single agent or in combination with immunotherapy.

Despite robust TET2 expression in MB49, there was no anti-tumor activity with single agent high-dose AA and no potentiation of anti-PD1 efficacy with the combination. In the MC38 model, with very high TET2 expression, high-dose AA markedly enhanced the efficacy of anti-PD1 immunotherapy even without significant single-agent activity.

Our data indicate that while TET2 plays an important role in AA-mediated anti-tumor activity, other proteins for which AA serves as a co-factor very likely participate in the cumulative AA-induced anti-tumor response. The HIF pathway is another documented AA co-factor target and mediator of anti-tumor activity *in vivo.* Negative correlations between tumor AA levels and HIF activity have also been reported in patients with solid tumor malignancies ^26,27^. Our data in the A20 model is consistent with this, showing downregulation of HIF target genes in AA-treated tumors of the shNC group, but not the sh*Slc23a2* group. Interestingly, AA did not decrease HIF target genes in the AA-resistant Renca tumors, despite an increase in intratumoral AA concentration. TET2 has been recently shown to be involved in inhibiting HIF signaling in renal cell carcinoma^28^; it is therefore possible that the lack of TET2 expression in Renca underlies the lack of downregulation of HIF target genes with AA. Together, the data suggest that the HIF pathway may also be contributing to anti-tumor function of AA. Given that the Km of asparaginyl hydroxylase (FIH) for AA is higher than that of prolyl hydroxylase^29^, it is likely that HIF transcriptional activity and target gene expression are affected before HIF-1α degradation, with deficiency of AA. The concentrations of intratumoral ascorbic acid in our study were in the range that would influence the optimal activity of these enzymes (TET, PHD, FIH), which is known to be around 1-2mM AA for collagen prolyl-4-hydroxylase ^30–33^. These findings are consistent with a previous report of high-dose IV AA administration in human colon cancer patients just prior to surgery in which AA co-factor activity (inhibition of HIF pathway) was significantly correlated with tumor AA levels while no oxidative damage (measured by γH2AX) was observed in tumors ^12^. It is also possible that additional epigenetic factors such as Fe^2+^ and 2-OG-dependent histone lysine methyltransferases for which AA serves as a co-factor may also play a role in AA-induced immune recognition ^34,35^, as well as a recently described direct post-translational modification of lysine residues by AA - contributing to enhanced STAT-1 mediated interferon signaling and HLA class 1 expression ^36^. Furthermore, it is likely that AA’s direct effect on the immune and non-immune microenvironment via activation of specific Fe^2+^ and 2-OG-dependent dioxygenases may be playing an additional role in its anti-tumor efficacy. For example, we have previously demonstrated that CD8+ T cells directly exposed to AA demonstrate enhanced cytotoxic ability against lymphoma cells in a coculture system, associated with CD8+T cell TET activation and increase in global 5-hmC levels ^3^. As such, a multitude of mechanisms involving specific cofactor activity across different cell types within the tumor contribute to AA’s overall anti-tumor effects, including the enhancement of anti-PD1 immunotherapy.

With regard to ascorbate transporters, most mammalian cells express SVCT2, while SVCT1 is expressed primarily on epithelial cells such as kidney, intestine, and liver. Both SVCTs carry out sodium-dependent transport of ascorbate; however functional experiments from cloned SVCT1/SVCT2 proteins have indicated that SVCT2 is high affinity/low capacity and SVCT1 is high capacity/low affinity^37^. In breast cancer cell line models, SVCT2 expression has previously been shown to be required for AA uptake *in vitro* and required for the AA anti-tumor response *in vivo* ^38^. This is consistent with our observation in the A20 model. The significant decrease in tumor ascorbate observed with *Slc23a2* knockdown suggests SVCT2 to be the primary uptake mechanism in A20 cells. Knockdown of *Slc23a2* in A20 prevented robust 5-hmC induction and also prevented the downregulation of HIF target genes with high-dose AA parenteral AA.

Our data clearly demonstrate that parenteral administration of ascorbic acid and supra-physiologic levels can further enhance the intratumoral activity of 2OG dependent enzymes including TET, PHD, FIH over physiologic ascorbic acid concentrations in a Gulo intact mouse model (which when extrapolated to humans would strongly indicate that intravenous administration would have a higher effect on these enzymes intratumorally compared to oral administration, given the tightly regulated gut absorption via SVCT1.) A few potential limiting factors to ascorbate uptake in cancers include tissue characteristics such as high cell density, poor perfusion, and reduced expression/activity of transporters ^1^. Our data are consistent with findings from a three-dimensional pharmacokinetic model to measure ascorbate diffusion suggesting that ascorbate penetration and distribution in a poorly vascularized tumor may require higher than normal plasma concentrations ^39^.

### Lack of oxidative stress with high-dose parenteral ascorbic acid *in vivo*

This study demonstrates that high-dose parenteral ascorbic acid (AA) treatment does not induce intratumoral oxidative damage *in vivo*, in either the sensitive A20 model or the resistant Renca model. Instead, AA treatment increases total antioxidant capacity within the tumors. In contrast to *in vivo* models, we show that oxidative damage is induced readily *in vitro*, highlighting a remarkable difference between *in vitro* and *in vivo* studies with high-dose AA. AA millimolar concentrations that did not cause any oxidative damage intratumorally *in vivo*, caused robust oxidative damage *in vitro*. Furthermore, if oxidative stress with generation of H_2_O_2_ in the tumor interstitial space was an important anti-tumor mechanism *in vivo*, knockdown of *Slc23a2* would not have fully reversed the anti-tumor effect of high-dose ascorbate, as H_2_O_2_ would be expected to diffuse across cell membranes without the necessity of a transporter. Finally, high dose parenteral AA did not have any single agent activity and did not potentiate anti-PD1 in the MB49 model despite its very low catalase expression, providing further evidence against the H_2_O_2_ mechanism *in vivo*. Because oxidative stress is generally considered a primary anti-tumor mechanism of parenteral high-dose AA, largely based on an abundance of literature with indisputable *in vitro* oxidative effects with high-dose AA and extrapolation from AA pharmacokinetics, these findings represent a major advance in our understanding of the mechanism of anti-cancer activity of high-dose AA *in vivo*.

We then hypothesized and demonstrated, using dynamic real-time extracellular H_2_O_2_ measurements with high-dose AA, that the difference in molecular oxygen concentration between standard *in vitro* and hypoxic *in vivo* conditions is an important factor underlying the marked discrepancy between the abundant *in vitro* and absent *in vivo* intratumoral oxidative stress with high-dose AA. The higher cell density along with a dense/clustered architecture of tumors also likely confer an enhanced ability to neutralize *any* extracellular H_2_O_2_ compared to cancer cells *in vitro*.

Doxorubicin, an agent which is well known to cause oxidative stress, did result in an increase in oxidative stress markers in A20 tumors. This indicates that the discrepancy between *in vitro* and *in vivo* oxidative stress does not extend to all pro-oxidant therapeutic agents and provides a positive control for intra-tumoral detection of oxidative damage. Combined treatment of doxorubicin and AA did not affect the response to doxorubicin and prevented DOX-induced generation of the oxidative marker 8-OHdG. These data suggest that doxorubicin’s primary mechanism of action is not oxidative damage, and that AA administered concurrently with doxorubicin does not exacerbate oxidative damage.

While we had previously considered the oxidative hemolysis reported after high-dose AA administration in G6PD patients to be evidence of *in vivo* oxidative effects of AA ^1,40,41^, we now propose that these patients may be uniquely susceptible to AA-induced oxidative damage ^42^. Our rationale follows that the absence of SVCT transporter expression/ascorbate uptake in red blood cells and corresponding reliance on uptake of dehydroascorbic acid (DHA) via glucose transporters results in DHA-induced oxidative damage (not H_2_O_2_-mediated oxidative damage), in the context of G6PD deficiency/impaired antioxidant machinery with insufficient NADPH^+^ and reduced glutathione.

It is possible that in the study by Chen et al.^11^, the exposure to atmospheric molecular oxygen concentration during extraction and processing may have influenced the amount of H_2_O_2_ produced and measured in the probe eluate from the tumor interstitial space. The authors reported that a steady state concentration of ⁓150uM H_2_O_2_ was generated in the tumor interstitial space (30min to 3hr period was reported) after intraperitoneal administration of a single dose of 4g/kg AA. However, our cumulative data strongly suggest that following parenteral AA administration, both the rate and amount of H_2_O_2_ production *in vivo* in a hypoxic tumor microenvironment with a tight cluster of cancer and stromal cells, is very likely much lower than previously thought. Furthermore, the higher cell density in tumors (and tumoral architecture) *in vivo* compared to cells *in vitro* confers a higher ability to counter H_2_O_2_-mediated oxidative effects. These factors underlie the absence of intratumoral oxidative damage in our study with sensitive and resistant models with the same intraperitoneal AA dose (4g/kg) daily, as well as in a study with intravenous AA administration in human colon cancer patients^12^. This contrasts with the consistent *in vivo* demonstration of increased intratumoral antioxidant/cofactor activity across multiple studies with parenteral high-dose ascorbate^2,3,5,12,20,43^. It is possible that some tumors with a high intracellular labile iron pool might trigger an intracellular prooxidant effect with increase in ascorbate content, but the data suggest that this is not a common feature with parenteral high-dose AA administration.

In summary, this study demonstrates that enhanced specific antioxidant/cofactor activity, not oxidative damage, is primarily responsible for the *in vivo* anti-cancer activity of high-dose parenteral AA including potentiation of anti-PD1 immunotherapy - with TET2 expression likely being necessary but certainly not sufficient. It demonstrates a marked discrepancy between the abundant *in vitro* and absent *in vivo* oxidative damage by high-dose AA, and elucidates the biochemical mechanisms underlying the discrepancy.

### Key implications of findings

Apart from representing a paradigm shift in the understanding of the cumulative anti-cancer mechanisms of high-dose ascorbic acid, these data have critical implications not just for the clinical translation of AA in cancer treatment (including in enhancing immunotherapy efficacy), but also the field of free radical biology:

First, for patients undergoing anti-PD1 immunotherapy treatment, could there be a potential benefit in high-dose ‘oral’ ascorbic acid particularly in those with deficient plasma levels of AA, despite the highly regulated gut absorption, to increase the intratumoral activity of 2OG dependent enzymes such as TETs, PHDs, FIH? Our data suggests so, and clinical studies are likely warranted to study this possibility, particularly considering the immense roadblocks in studying the much more promising intravenous high-dose ascorbic acid in high-quality/randomized clinical trials. However, it is important to note that the AA-sensitive A20 syngeneic model had an intact Gulo enzyme and was therefore not ‘deficient’ in plasma AA concentration; yet the administration of parenteral high-dose AA in this model resulting in supra-physiologic plasma and intratumoral AA concentrations demonstrated synergy with anti-PD1^3^. Together, the results strongly suggest that for humans, the intravenous route would be the ideal mode of administration even though the mechanism of its anti-cancer effect is mediated primarily by cofactor activity. A 3-arm randomized trial with no, oral, or IV AA (in combination with anti-PD1 immunotherapy) with biomarker assessment would be the definitive method to study the incremental benefit of IV AA over oral AA and without supplementation, in enhancing the efficacy of immunotherapy.

Second, given the marked discrepancy between H_2_O_2_ generation and viability in a standard cell culture incubator (with supraphysiologic O_2_ concentration) and in a hypoxia incubator (with O_2_ levels representative of mammalian tissue O_2_ concentration), lab-based free radical biology experiments conducted with standard cell culture incubators, including the interpretation of multiple signaling pathways affected by free radicals, need to be examined for consistency *in vivo* prior to determining relevance to human physiology.

## Methods

### Cell culture

A20, Renca, and 786-O cell lines were purchased from ATCC. MB49 mouse urothelial carcinoma line was purchased from AddexBio Technologies (San Diego, CA). MC38 mouse colon adenocarcinoma line was purchased from Kerafast (Shirley, MA). Renca was cultured in RPMI or DMEM supplemented with 10% fetal bovine serum (FBS, Gemini Benchmark). 786-O and A20 were cultured in RPMI 1640 with 10% FBS. MB49 was cultured in DMEM with 10% FBS. MC38 was cultured in DMEM with 10% FBS, 2 mM glutamine, 0.1 mM nonessential amino acids, and 1 mM sodium pyruvate. All cell lines were cultured in the presence of 100 U/ml penicillin and 100 µg/ml streptomycin (Gemini) and maintained at 37° C in 5% CO_2_.

### Cell treatment

Cell lines were treated with specified concentrations and timepoints of L-ascorbic acid (Fisher Scientific), with or without catalase (Sigma). For catalase treatment, cells were pre-treated for 30 min with catalase prior to L-ascorbic acid treatment. Cells were treated with specified concentrations of H_2_O_2_ (Sigma) as a positive control.

### shRNA and ORF expression plasmids

A20 cells were transduced with MISSION TRC1.5 or TRC2 PLKO.5-PURO shRNA lentivirus constructs (Sigma) targeting murine*Tet2* (Sigma TRCN0000250894), *Slc23a2* (Sigma TRCN0000070162) or non-targeting control (Sigma SHC216). Briefly, lentivirus was added to A20 cells at MOI=5-10 and transduced by spinoculation (90 minutes, 900-1000 x g, room temp) followed by an additional 3 h incubation with lentivirus at 37 C. Renca cells were transduced with pReceiver-Lv105 lentivirus empty vector (EX-NEG-Lv105, GeneCopoeia) or ORF expression clone for murine *Tet2* (NM_001040400.2, GeneCopoeia) by applying filtered lentivirus supernatant directly to the cells. All transduced cells were selected at 48 h post-transduction using puromycin, and knockdown efficiency was assessed by Western blot and/or qPCR. Renca single cell clones were generated by limiting dilution to ensure a similar level of expression across cells. Transduced cell lines were maintained in 0.25 μg/ml puromycin. Puromycin was removed from culture media 48-72 h prior to use in mouse tumor injections.

### Cell viability assay

Relative cell growth was compared *in vitro* using the Cell Titer Blue kit (Promega). For stable transduced cell lines, equal numbers of cells were plated per well for A20 (10,000 cells) and Renca (5,000 cells) and incubated for 72 hours. Cell growth was determined by the addition of Cell Titer Blue assay reagent and quantified on a plate reader (Cytation 3, BioTek). For treated cells, Renca (5,000 cells) and A20 (10,000 cells) were plated and incubated overnight. Cells were then treated with the specified concentrations of AA, H_2_O_2_ and/or catalase and incubated for 2 hours (A20) or 4 hours (Renca). Treatment media was removed, and cells were washed prior to incubation with Cell Titer reagent for 2 hours followed by quantification on a plate reader. For the cell density experiments (Suppl Fig S6, S7), cells were plated at different numbers as indicated, and the Cell Titer Blue assay was used as above.

### Mouse models

All mouse experiments were approved by the Albert Einstein College of Medicine and Northwestern University Institutional Animal Care and Use Committees. BALB/c and C57Bl/6 female or male mice (5-8 weeks old) were purchased from The Jackson Laboratory (Bar Harbor, ME) or Charles River. After acclimation, female BALB/c mice received a subcutaneous flank injection of 5x10^6^ A20 or 5x10^5^ RENCA tumor cells, and C57Bl/c mice (MB49: male; MC38: female) received 5x10^5^ MB49 or MC38 tumor cells in PBS. Once tumors were palpable, mice were randomly assigned to treatment groups, and treatment was initiated (AA studies (Figs 1-3): n≥5 mice per treatment group; Doxorubicin study (Fig 10): n=4 per group; AA and Doxorubicin study (Fig 8): n≥7 mice per group; AA and anti-PD1 studies (Fig 1,9, Suppl Fig 8): n≥5 mice per treatment group). Mice in which palpable tumors were not observed at the initiation of treatment were excluded from the study. Mice receiving AA were given daily intraperitoneal (IP) injections of sodium L-ascorbate (Sigma-Aldrich, St. Louis, MO). To balance the osmotic effect of sodium L-ascorbate, mice in the Vehicle group were administered NaCl (Sigma-Aldrich). To allow mice to acclimate to the osmotic load, 1.5 M ascorbate or NaCl injection volumes were gradually increased over the course of 4 days as previously described ^3^. After this, 300 μl injections of 1.5 M ascorbate or NaCl were given daily. For the studies with combined AA and anti-PD1 treatment, AA or NaCl was given daily, and 200 µg anti-PD1 (BE0146; BioXCell) or isotype control (BE0089; BioXCell) was given every other day as previously described^3^. In the Doxorubicin (Dox) only treatment study, mice were administered a single intraperitoneal (IP) injection of 1 mg/kg doxorubicin hydrochloride (Pfizer) diluted in saline or vehicle (Veh) every 3 days, resulting in a cumulative dose of 4mg/kg. In the Doxorubicin (Dox) + AA treatment study, mice received a single IP injection of 1 mg/kg doxorubicin hydrochloride (Pfizer) diluted in saline or vehicle (Veh) every 2 days, resulting in a cumulative dose of 5mg/kg. For all studies, tumor volume (mm^3^) was monitored every one to two days over the duration of the study by caliper as previously described ^3^. Treatment continued until the first mouse reached tumor size or other humane endpoint criteria. Mice were sacrificed by CO_2_ and tumors were removed, weighed, and divided for downstream assays. Tumor sections were fixed in formalin, frozen, and processed fresh for ascorbate measurement (shRNA and overexpression vector studies only).

### Nucleic acid extraction

DNA was isolated from frozen cells and tumor tissue using Qiagen DNeasy Blood and Tissue kit. RNA was isolated from frozen cells and tumor tissue using Qiagen RNeasy kit.

### PCR

cDNA was reverse transcribed from RNA using SuperScript III (Invitrogen) or High-Capacity cDNA Reverse Transcription kit (Applied Biosystems). For analysis by gel electrophoresis, PCR was performed using HotStarTaq (Qiagen). Primers sequences for mouse *Slc23a1* (SVCT1), and *Actb* (β–Actin) were previously published ^44^. The following primers were used for mouse *Slc23a2*, Forward/Reverse: ACAGAGAACGGCATTGCAGA/TGGCCTGAAACAGGGGTAAC. PCR products were visualized by agarose gel electrophoresis. For quantitative PCR (qPCR), the following mouse primer sets were used: *Tet2* ^45^ Forward/Reverse: TGCTTTCCCAACACGGAACTA/GCACCATTAGGCATTAGCACAAT; *Slc23a2* (Origene) Forward/Reverse: GGACGGCATACAAGTTCCAGCT/AGCCATAGTCGGTGCTGTTGGA; *Vegfa* Forward/Reverse: CTGCTGTAACGATGAAGCCCTG/GCTGTAGGAAGCTCATCTCTCC; *Car9* Forward/Reverse: GTGCACCTCAGTACTGCTTT/CTTCCTCCGAGATTTCTTCCA; *Glut1 (Slc2a1)* Forward/Reverse: GCTTCTCCAACTGGACCTCAAAC/ACGAGGAGCACCGTGAAGATGA; *Actb* Forward/Reverse: CATTGCTGACAGGATGCAGAAGG/TGCTGGAAGGTGGACAGTGAGG; and *Hmbs* (Origene) Forward/Reverse: GTGCCTACCATACTACCTCCTG/ACTCTCCTCAGAGAGCTGGTTC. Relative transcript expression was quantified on QuantStudio Real-Time PCR System (Applied Biosystems) instrument with PowerUp SYBR Green Master Mix (Applied Biosystems) and CFX Biorad instrument with SsoAdvanced Universal SYBR® green, both using the delta delta ct method. *Actb* and *Hmbs* were used as housekeeper genes for cell line and tumor tissue, respectively.

### 8-OhdG ELISA

8-OHdG was quantified using EpiQuik 8-OHdG DNA Damage Quantification Direct Kit (Epigentek). An input DNA amount of 200-300 ng was used for each tumor. The assay was performed following manufacturer’s instructions. Standards were used to calculate ng 8-OHdG detected per ng input DNA (8-OHdG%).

### 5-hmC dot blot

DNA dot blot for 5-hmC was performed as previously described ^19^. Cell line DNA was denatured by incubation at 95 C for 10 min followed by neutralization. DNA was then spotted onto a nylon membrane (Amersham Hybond N+) and baked for 30 min at 80 C. The membrane was incubated in anti-5-hmC (Active Motif cat no. 39069) overnight at 4 C followed by secondary HRP-conjugated anti-rabbit antibody (Cell Signaling #7074) for 1 h at room temp. 5-hmC was visualized by chemiluminescence and detected by film or Odyssey XF (LI-COR). Methylene blue (0.04%) was used as a loading control.

### Ascorbic acid and antioxidant capacity measurement

Tumor ascorbic acid content and antioxidant capacity was measured by Abcam’s Ascorbic Acid Assay Kit (ab65656) following manufacturer’s instructions. Briefly, approximately 10 mg fresh tumor tissue was homogenized, and antioxidant capacity was measured by reduction of Fe^3+^ to Fe^2+^ and detected by colorimetric probe (absorbance reading at 545-600 nm). The assay was performed in parallel samples with one containing ascorbate oxidase for ascorbate removal allowing for measurement of background antioxidant capacity. Ascorbate content was determined by subtraction of background antioxidant capacity from total antioxidant capacity and calculated using a standard curve. Ascorbate was normalized to total protein and expressed as nmol ascorbate per mg protein. Total antioxidant capacity was quantified as the absorbance in wells without ascorbate oxidase per mg protein.

### Western blot

Lysates from cell line or flash-frozen tissue samples were prepared using RIPA lysis buffer (Cell Signaling). Primary antibodies against SVCT1 (Abcam, AB236878), TET2 (Cell Signaling Technology, 18950), TET2 (mouse-specific, Cell Signaling, 92529), TET1 (GeneTex [N3C1]), TET3 (GeneTex [C3]), catalase (D5N7V, Cell Signaling, 14097) and β-Actin (Novus Biologicals, NB600-501) recognized both human and mouse proteins for the respective targets unless noted as species-specific. Equal amounts of each lysate (30-40 µg) were loaded. Proteins were detected using IR-Dye secondary antibodies (LI-COR) on Odyssey FC or Odyssey XF instruments (LI-COR). Quantification was performed using LI-COR Image Studio and Empiria software and proteins of interest were normalized to total protein for protein abundance.

### Immunofluorescence

Immunofluorescence for 8-OhdG and 4-HNE was performed on cell line and tissue samples. For A20, cells were plated onto coated coverslips (A20 cells: 0.24 x10^6^ cells/well in 24 well plate) and allowed to adhere for 1h. Renca cells were plated on coverslips (0.5x10^5^cells per well of 6-well plate) and allowed to adhere overnight. Cells were then treated with H_2_O_2_ for 4 or 6 h and coverslips were washed with PBS and fixed with formalin for 10 min. Fixed cells were incubated with 0.1 M glycine for 10 min at room temp. Cover slips were incubated in blocking buffer, followed by primary antibody (4-HNE 1:50, Abcam ab46545; 8-OxO-dG 1:250, Trevigen clone 15A3) and secondary antibody (anti-rabbit IgG Alexa fluor-488 and anti-mouse IgG Alexa Fluor-568; Thermo Scientific), each for 1h at room temp. Coverslips were mounted in SlowFadeTM Gold antifade reagent with DAPI. For tumors, fresh tumor sections were placed in formalin for 48 h after which time they were transferred to 70% ethanol prior to paraffin embedding. Slides with 4 µm FFPE sections were rehydrated followed by antigen retrieval with Universal HIER antigen retrieval reagent (Abcam). Tissues were then incubated with blocking buffer for 30 min at room temp, primary antibody against 4-HNE (1:100) and 8-OxO-dG (1:100) overnight at 4C, then secondary antibody anti-rabbit IgG Alexa fluor-488 and anti-mouse IgG Alexa Fluor-568 at room temperature for 1h. Coverslips were mounted in SlowFadeTM Gold antifade reagent with DAPI. Stained slides were visualized, and images captured at 40x on an Olympus CKX53 fluorescent microscope. The image was merged using ImageJ software.

### Immunohistochemistry and scoring

Immunohistochemistry for CD8 was performed by Northwestern University Mouse Histology & Phenotyping Laboratory. Formalin fixed paraffin-embedded tumors were stained for CD8a (eBioscience cat# 14-0195-82; 1:150 dilution) and detected with Vectastain ABC HRP kit (Vector Labs). Three to five representative 400 µm^2^ regions were scored for number of CD8-positive cells. These scores were averaged for each tissue.

### Malondialdehyde (MDA) assay

Lipid peroxidation was measured by MDA assay kit (Abcam, ab233471). Tissue extracts were prepared from flash frozen tumor using 20 mM sodium phosphate buffer + 0.5% TritonX-100, pH 3. Lysates were analyzed for MDA abundance following manufacturer’s instructions. Total protein was determined in lysates and data were reported as ng MDA per µg protein.

### Dynamic real-time extracellular H_2_O_2_ detection

Equal numbers of Renca (5,000) cells were plated per well of a 96-well plate and incubated overnight before treatments. BIOTRACKER green free H_2_O_2_ dye (Millipore Sigma, cat # SCT040) was added at a final concentration of 1µM and cells were allowed to equilibrate to hypoxia (3% O_2_) or normoxia (19% O_2_) conditions.

Cells were then treated with 1mM or 5mM of ascorbic acid (or 0.1mM of H_2_O_2_ for the positive control) and immediately incubated for 10 hours under continuous hypoxia (3% O_2_) or normoxia (19% O_2_) conditions, both in the presence of 5% CO_2_. Serial real-time extracellular hydrogen peroxide measurements were taken over the course of 10 hours within the respective conditions, every 30 min, on a Cytation3 plate reader (BioTek) equipped with CO_2_/O_2_ and temperature control at the wavelength of 516 nm.

### Growth pattern/time-based differences in cytotoxicity and biochemical activity

Some adherent cell lines, such as Renca, preferentially grow in colonies/clusters, where the new daughter cells formed via mitosis remain adherent to the parent cell. In such cell lines, a longer pretreatment incubation time (with higher no. of cell divisions) gives a ‘clustered’ pattern, whereas a shorter incubation time (with lower no. of cell divisions) gives a ‘scattered’ pattern. Other adherent cell lines such as 786-O do not exhibit this ‘clustered’ pattern of growth over time, as the daughter cells largely detach from the parent cell after mitosis. For the experiment in Fig S7, a smaller number of seeded Renca cells with a longer incubation period was used to generate the ‘clustered’ pattern, and a larger number of seeded Renca cells with a smaller incubation period was used to generate the ‘scattered’ pattern, ensuring that the cell count at treatment was similar between the ‘clustered’ and ‘scattered’ groups (summarized in Figure S7A).

### Single nuclei RNA sequencing

Four representative tumors were selected from each MB49 group (Vehicle, AA, anti-PD, and AA+anti-PD1) (Fig 5). Nuclei were isolated from flash frozen tumor tissue using 10x Genomics Chromium Nuclei Isolation kit with RNase Inhibitor following manufacturer’s instructions. ProtectRNA RNase Inhibitor (Sigma-Aldrich) was included for all samples throughout processing. Nuclei were fixed using the Evercode Nuclei Fixation kit v3 (Parse Biosciences). Single nuclei suspensions were visually assessed by microscope for quality. Libraries were constructed in the NUSeq Core using the Evercode WT Mega v3 kit. Three sub-libraries containing ∼6,000 nuclei per sample were sequenced at Parse Biosciences. FASTQ files were processed using the Parse Biosciences Pipeline (v1.5.0, Parse Biosciences).

Trailmaker^TM^ (Parse Biosciences) was used to complete our single cell RNA-sequencing data analysis. Unfiltered count matrices were uploaded to Trailmaker, and background was removed by setting a minimum transcripts per cell threshold on a per sample basis (threshold range: 400 to 500). Dead or dying cells were removed by filtering barcodes with high mitochondrial content (3-7.5% cut-off). Outliers in the distribution of number of genes vs number of transcripts were removed by fitting a spline regression model (p-values between 0.0003 and 0.001). Cells with a high probability of being doublets were filtered out using the scDblFinder method (threshold range: 0.48 to 0.72). Data normalization, principal-component analysis (PCA) and data integration using Harmony were performed on data from high-quality cells. Clusters were identified using the Leiden method, and a Uniform Manifold Approximation and Projection (UMAP) embedding was calculated to visualize the results. Clusters were annotated for immune cells using ScType.

### Variant Analysis

To assess variants in the *Tet2* locus for the MB49 cell line, we accessed RNA-seq files from GSE112973 ^46^. Fastq files were processed using fastp and aligned using BWA. Variants were called using LoFreq and annotated using snpEff.

### Statistics

Data were first assessed by Kolmogorov-Smirnov test for normality and Grubbs’ outlier test. Outliers (p<0.05 in Grubbs’ outlier test) were removed from relevant analyses. Statistical comparisons between two groups were performed using Student’s or Welch’s T tests.

## Author contributions

RAL and NKS designed the experiments; TA, RAL, SS, VD, RA, HS, and NKS performed the experiments; RAL, VD, TA, SS, and NKS analyzed the data; RAL and NKS wrote the manuscript; NKS was responsible for conceptualization, direction, and funding.

## Acknowledgments

This work was supported by funding from the Lotte & John Hecht Foundation grant #4809 (N.S.), the American Cancer Society Research Scholar Grant RSG-20-137-01 TBG (N.S.), and the Northwestern University Division of Hematology/Oncology (N.S.). Dynamic real-time extracellular hydrogen peroxide detection was performed in the Analytical BioNanoTechnology Equipment Core Facility of the Simpson Querrey Institute for BioNanotechnology at Northwestern University (H.S. Acting Director). ANTEC receives partial support from the Soft and Hybrid Nanotechnology Experimental (SHyNE) Resource (NSF ECCS-2025633) and Feinberg School of Medicine, Northwestern University. Non-clinical research histopathology and molecular phenotyping services were provided by the Northwestern University Mouse Histology and Phenotyping Laboratory (MHPL) which is supported by NCI P30-CA060553 awarded to the Robert H Lurie Comprehensive Cancer Center.

## SUPPLEMENTAL MATERIAL/EXTENDED DATA

**Supplemental Figure S1.**
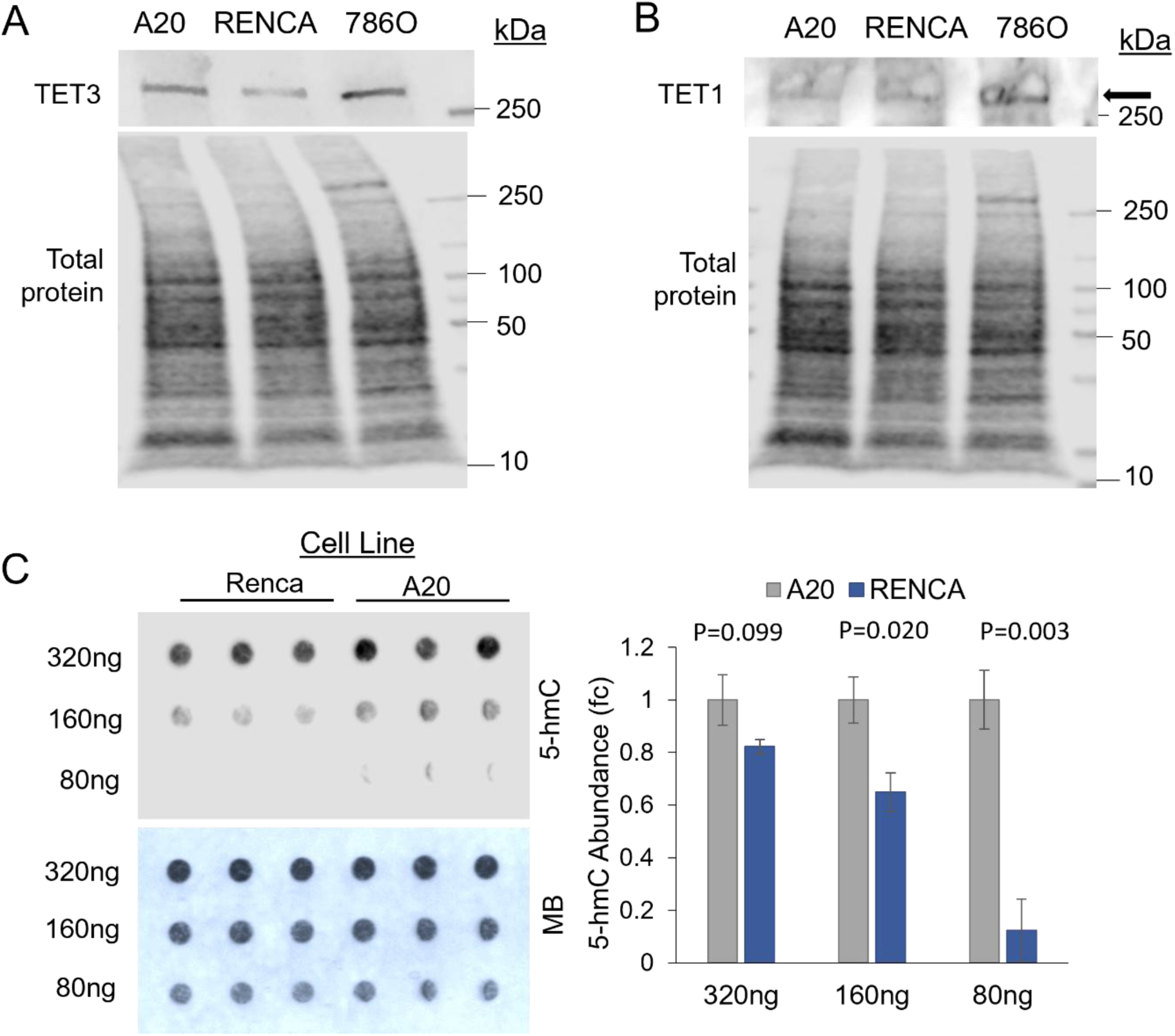
A,B. Protein expression of Tet enzymes across cell lines. C. 5-hmC in untreated Renca compared to A20 cells. 5-hmC, 5-hydroxymethylcytosine; MB, methylene blue loading control.

**Supplemental Figure S2.**
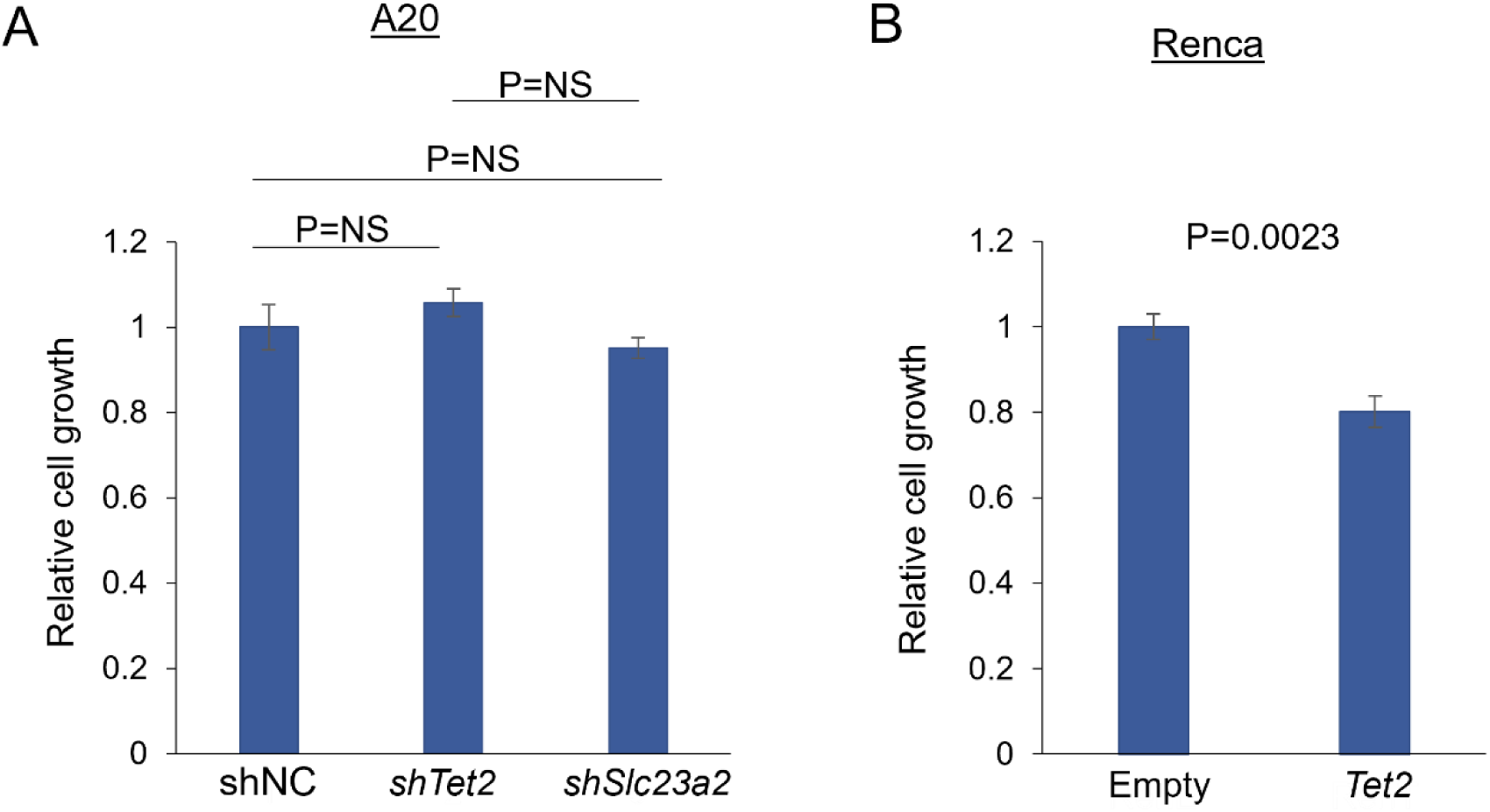
Cell Viability of stably selected A20 and Renca cells. Cell viability was assessed 72 h after plating equal numbers of each stably modified cell line. Data represented as mean ± se.

**Supplemental Figure S3.**
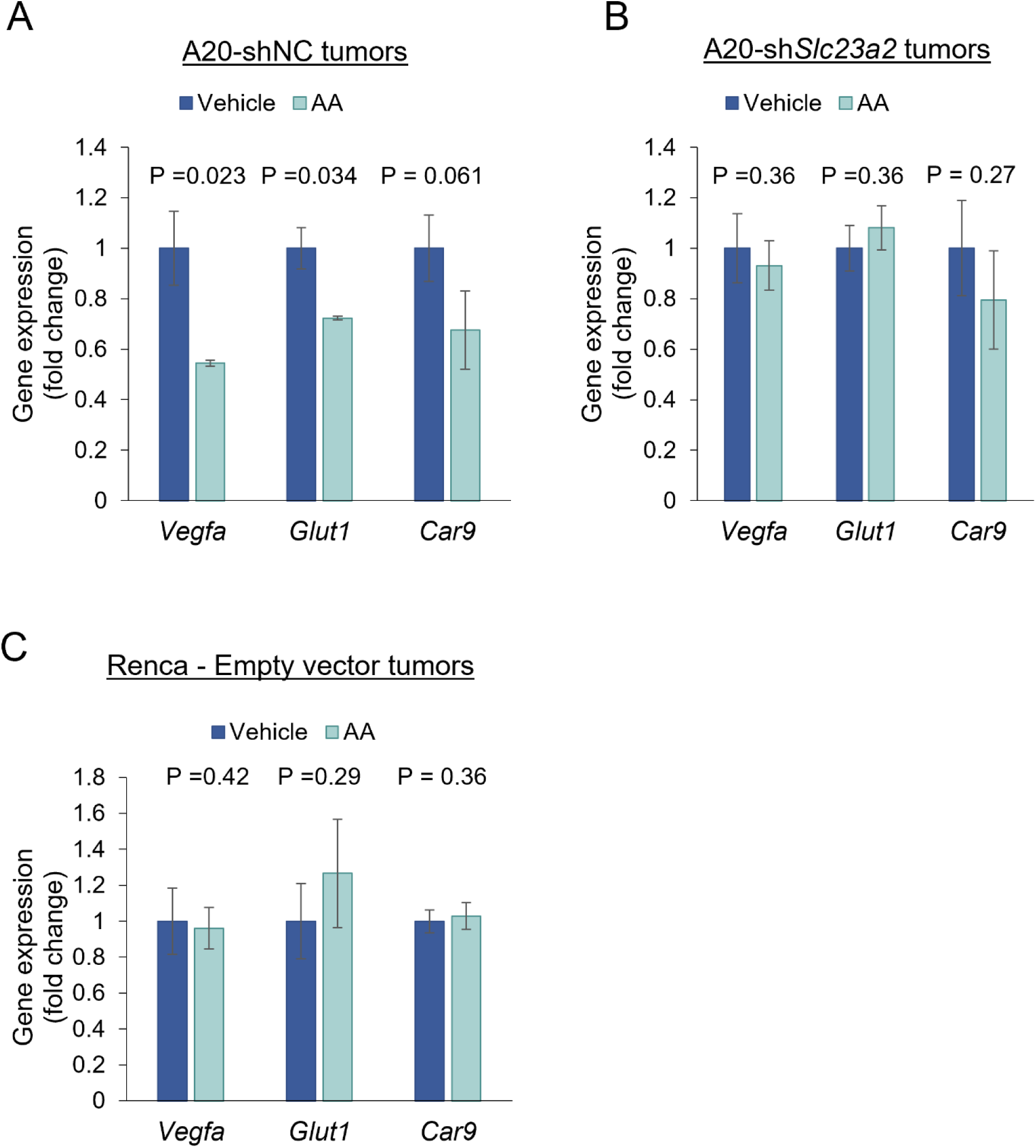
HIF pathway expression. (A) In the A20 model, HIF target gene expression is downregulated with AA treatment in (A) shNC but not in (B) sh*Slc23a2* tumors. (C) HIF target gene expression is not downregulated by AA in Renca tumors. Data represented as mean ± se.

**Supplemental Figure S4.**
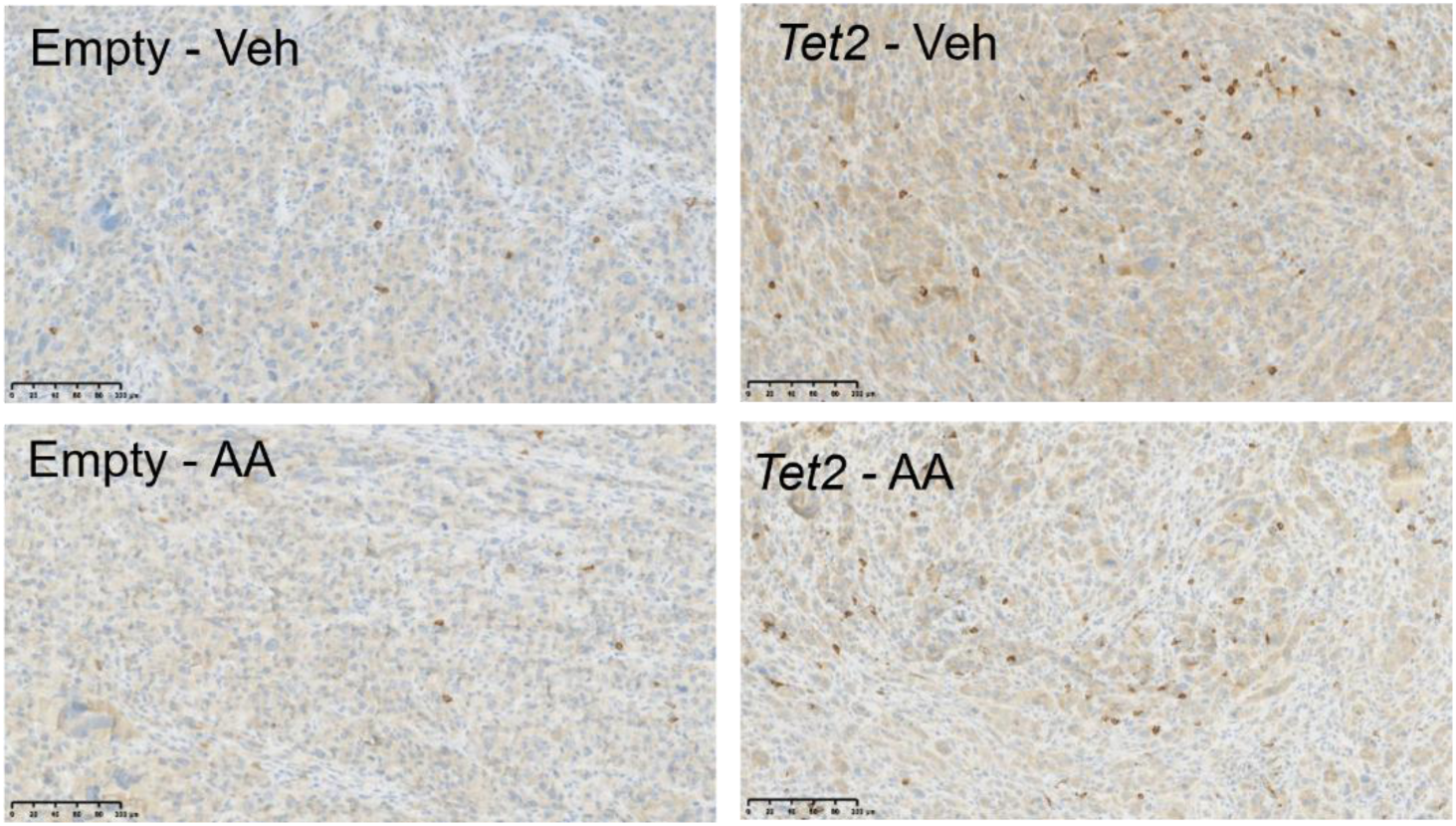
Renca CD8 Immunohistochemistry. Representative IHC CD8 stains for Renca-Empty and Renca-*Tet2* tumors.

**Supplemental Figure S5.**
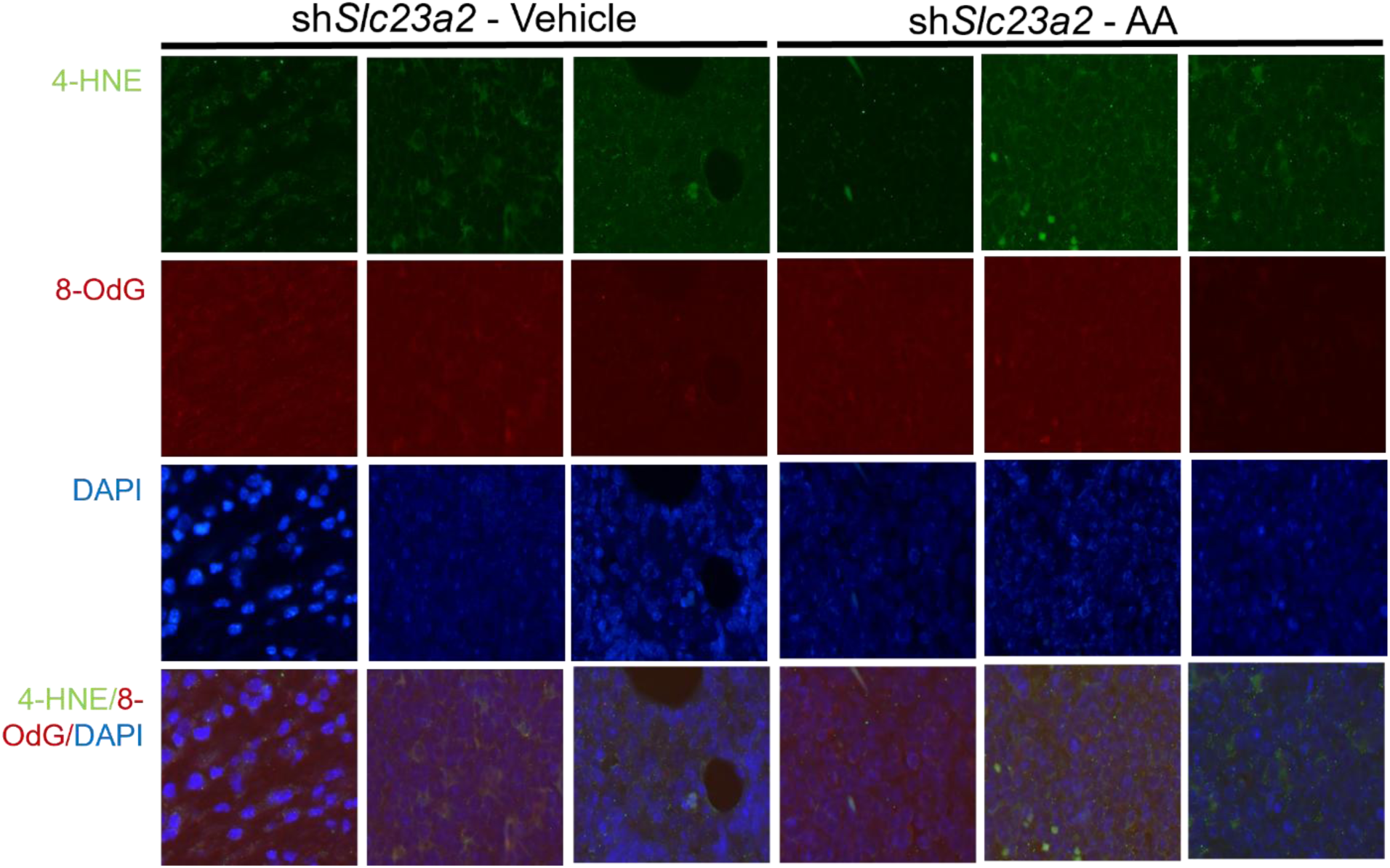
Knockdown of *Slc23a2* does not influence AA-induced oxidative stress in A20 model.

**Supplemental Figure S6.**
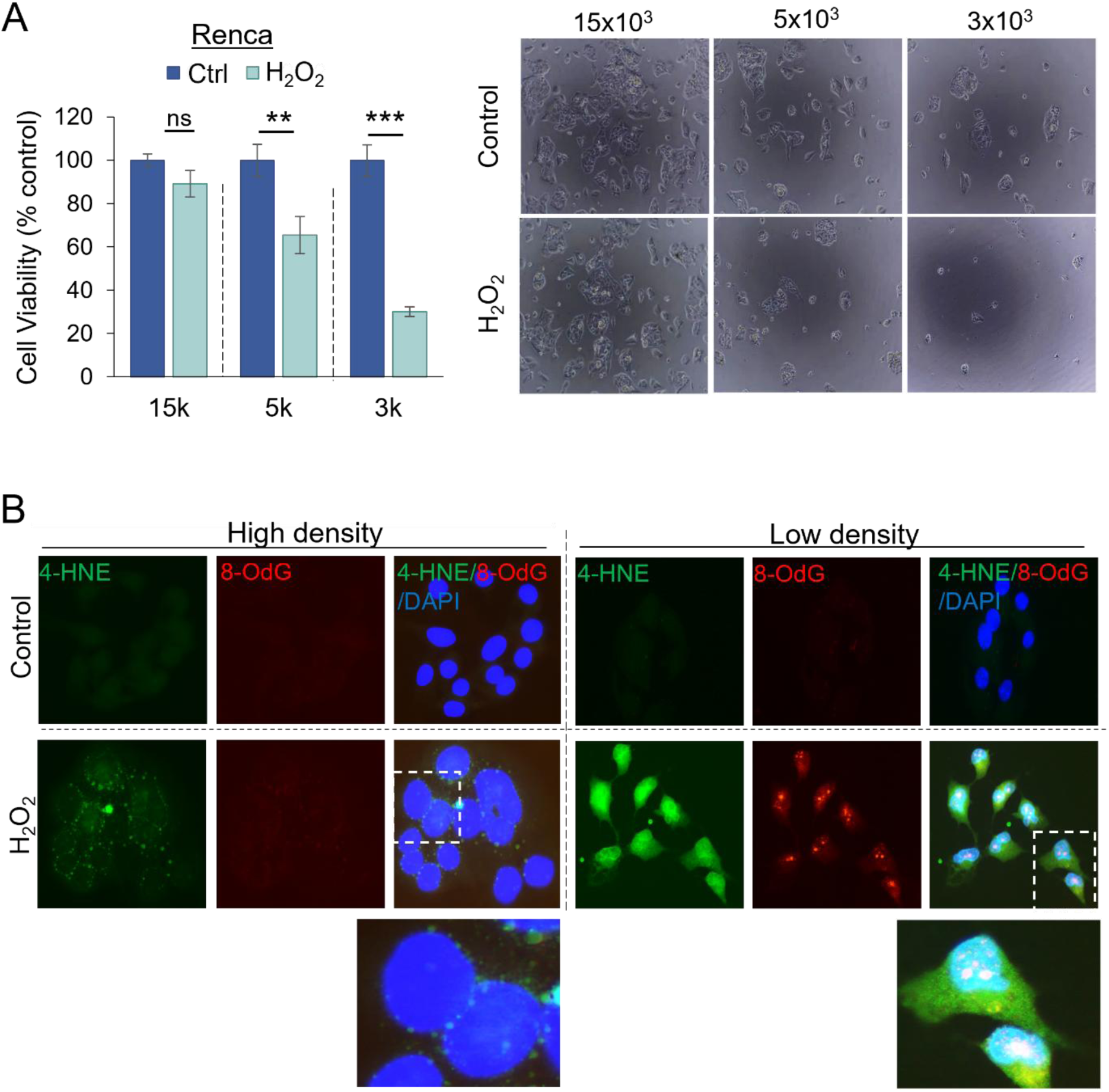
Impact of cell density on H_2_O_2_-induced oxidative stress and cytotoxicity. A. Cell viability and representative images of Renca cells treated with H_2_O_2_ (1mM for 2hr) with different pretreatment cell density as indicated (*: p<0.05; **: p<0.01; ***: p,0.001; 2-sided t-test). B. Immunofluorescence of oxidative stress markers 4-HNE and 8-Oxo-dG showing markedly higher staining with lower density compared with higher density (Renca; 1mM H_2_O_2_ for 2 hr). Data represented as mean ± se.

**Supplemental Figure S7.**
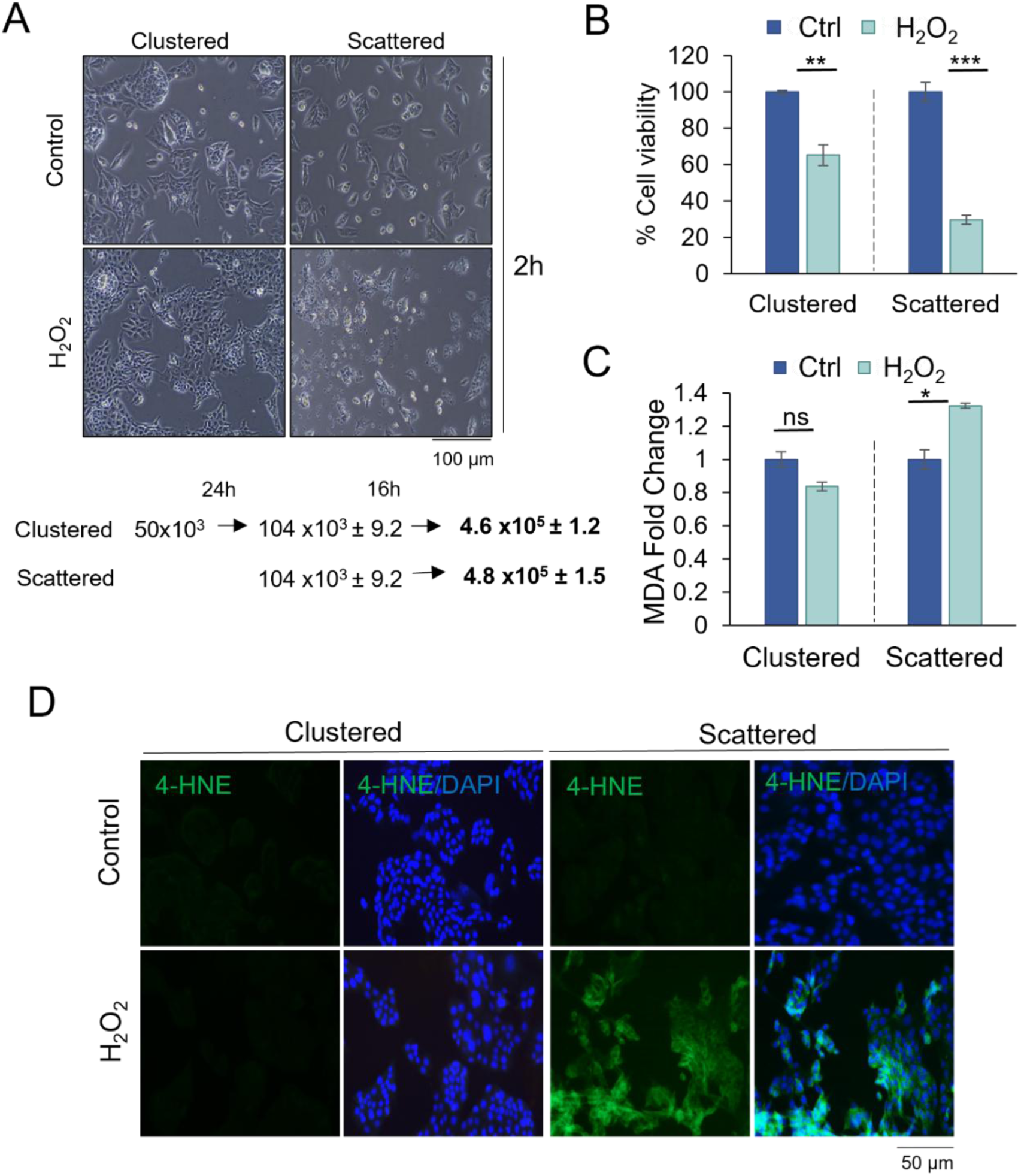
A. Representative images of Renca ‘clustered’ vs ‘scattered’ treated with H_2_O_2_ (1mM, 2h) vs control. Smaller number of seeded cells with a longer incubation period (’clustered’ pattern), and a larger number of seeded cells with a smaller incubation period (‘scattered’ pattern), ensuring that the cell count at treatment was similar between the ‘clustered’ and ‘scattered’ groups (Renca). B. With H_2_O_2_ treatment (1mM, 2h), there was marked reduction in viability after 24 h with ‘scattered’ compared to ‘clustered’ pattern in Renca cells, despite similar cell count at treatment and similar viability (absolute absorbance) of respective controls. C. Underlying the cytotoxicity difference, there was markedly higher oxidative damage with ‘scattered’ compared to ‘clustered’ pattern as measured by MDA (C) and 4-HNE IF (D) with H_2_O_2_ (1mM, 2h). Cells were fixed 24 h after treatment. Data represented as mean ± se.

**Supplemental Figure S8.**
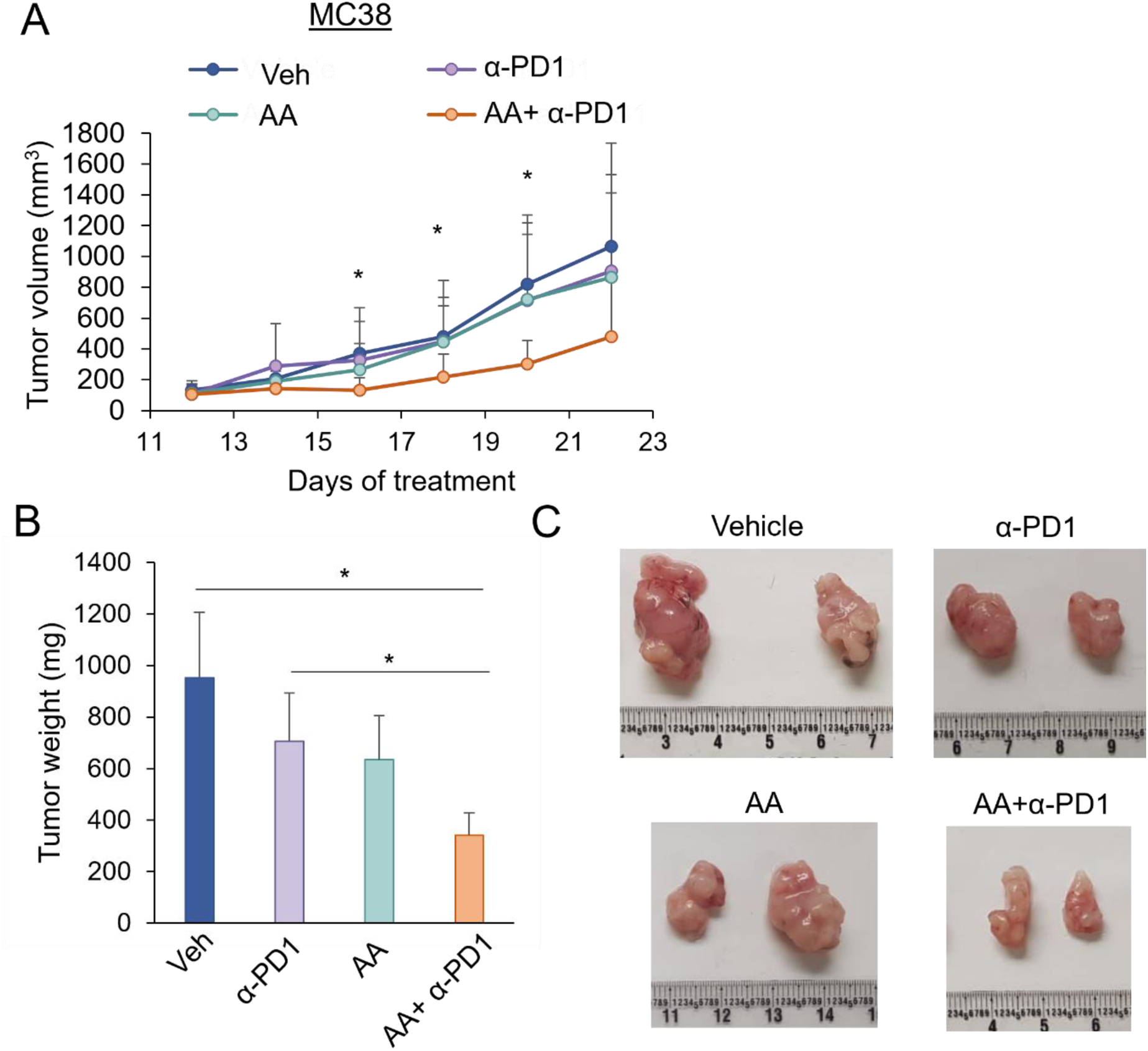
AA and anti-PD1 treatment in MC38 syngeneic model. A. Tumor volume over time in response to Vehicle (n=5), AA (n=6), anti-PD1 (n=7), and AA+anti-PD1 (n=7) treatment (mean ± sd). B. Final tumor weight (mean ± se) and (C) representative tumor images. P-values represent Welch’s T-test. *, p<0.05.

